# Clear: Composition of Likelihoods for Evolve And Resequence Experiments

**DOI:** 10.1101/080085

**Authors:** Arya Iranmehr, Ali Akbari, Christian Schlötterer, Vineet Bafna

## Abstract

The advent of next generation sequencing technologies has made whole-genome and whole-population sampling possible, even for eukaryotes with large genomes. With this development, experimental evolution studies can be designed to observe molecular evolution “in-action” via Evolve-and-Resequence (E&R) experiments. Among other applications, E&R studies can be used to locate the genes and variants responsible for genetic adaptation. Existing literature on time-series data analysis often assumes large population size, accurate allele frequency estimates, and wide time spans. These assumptions do not hold in many E&R studies.

In this article, we propose a method-Composition of Likelihoods for Evolve-And-Resequence experiments (Clear)–to identify signatures of selection in small population E&R experiments. Clear takes whole-genome sequence of pool of individuals (pool-seq) as input, and properly addresses heterogeneous ascertainment bias resulting from uneven coverage. Clear also provides unbiased estimates of model parameters, including population size, selection strength and dominance, while being computationally efficient. Extensive simulations show that Clear achieves higher power in detecting and localizing selection over a wide range of parameters, and is robust to variation of coverage. We applied Clear statistic to multiple E&R experiments, including, data from a study of *D. melanogaster* adaptation to alternating temperatures and a study of outcrossing yeast populations, and identified multiple regions under selection with genome-wide significance.

## 1 Introduction

Natural selection is a key force in evolution, and a mechanism by which populations can adapt to external ‘selection’ pressure. Examples of adaptation abound in the natural world [22], including for example, classic examples like lactose tolerance in Northern Europeans [9], human adaptation to high altitudes [55, 69], but also drug resistance in pests [15], HIV [24], cancer [27, 70], malarial parasite [3, 44], and others [56]. In these examples, understanding the genetic basis of adaptation can provide valuable information, underscoring the importance of the problem.

Experimental evolution refers to the study of the evolutionary processes of a model organism in a controlled [7, 10, 28, 37, 38, 46, 47] or natural [5, 8, 16, 17, 41, 50, 68] environment. Recent advances in whole genome sequencing have enabled us to sequence populations at a reasonable cost, even for large genomes. Perhaps more important for experimental evolution studies, we can now evolve and resequence (E&R) multiple replicates of a population to obtain *longitudinal time-series data,* in order to investigate the dynamics of evolution at molecular level. Although constraints such as small sizes, limited timescales, and oversimplified laboratory environments may limit the interpretation of E&R results, these studies are increasingly being used to test a wide range of hypotheses [34] and have been shown to be more predictive than static data analysis [12, 18, 52]. In particular, longitudinal E&R data is being used to estimate model parameters including population size [33, 49, 60, 64, 65, 67], strength of selection [11, 29, 30, 40, 43, 57, 60], allele age [40] recombination rate [60], mutation rate [6, 60], quantitative trait loci [4] and for tests of neutrality hypotheses [8, 13, 23, 60].

While many E&R study designs are being used [6, 53], we restrict our attention to the adaptive evolution due to standing variation in fixed size populations. This regime has been considered earlier, typically with *D. melanogaster* as the model organism of choice, to identify adaptive genes in longevity and aging [13, 51] (600 generations), courtship song [63] (100 generations), hypoxia tolerance [71] (200 generations), adaptation to new laboratory environments [26, 46] (59 generations), egg size [32] (40 generations), C virus resistance [42] (20 generations), and dark-fly [31] (49 generations).

The task of identifying selection signatures can be addressed at different levels of specificity. At the coarsest level, identification could simply refer to deciding whether some genomic region (or a gene) is under selection or not. In the following, we refer to this task as *detection.* In contrast, the task of *site-identification* corresponds to the process of finding the favored mutation/allele at nucleotide level. Finally, *estimation of model parameters,* such as strength of selection and dominance at the site, can provide a comprehensive description of the selection process.

In the effort to analyze E&R selection experiments, many authors chose to adapt existing tests that were originally used for static data, pairwise comparisons (two time-points) and single replicates to perform a null scan. For instance, Zhu *et al.* [71] used the ratio of the estimated population size of case and control populations to compute test statistic for each genomic region. Burke *et al.* [13] applied Fisher exact test to the last observation of data on case and control populations. Orozco-terWengel *et al.* [46] used the Cochran-Mantel-Haenszel (CMH) test [1] to detect SNPs whose read counts change consistently across all replicates of two time-point data. Turner *et al.* [63] proposed the diffStat statistic to test whether the change in allele frequencies of two populations deviate from the distribution of change in allele frequencies of two drifting populations. Bergland *et al.* [8] calculated *F_st_* to populations throughout time to signify their differentiation from ancestral (two time-point data) as well as geographically different populations. Jha *et al.* [32] computed test statistic of generalized linear-mixed model directly from read counts.

Alternatively, *direct* methods have been developed to analyze time-series data by taking a likelihood approach, and estimating population genetics parameters. Bollback *et al.* [11] proposed a Hidden Markov Model (HMM) to estimate the selection coefficient *s* and population size by using a diffusion approximation to the continuous Wright Fisher Markov process. Steinrücken and Song [57] proposed a general diploid selection model which takes into account of dominance of the favored allele and approximates likelihood analytically. Mathieson and McVean [43] adopted HMMs to structured populations and estimated parameters using an Expectation Maximization (EM) procedure on discretized allele frequency. Feder *et al.* [23] modeled increments in allele frequency with a Brownian motion process, proposed the Frequency Increment Test (FIT). More recently, Topa *et al.* [62] proposed a Gaussian Process (GP) for modeling single-locus time-series pool-seq data. Terhorst *et al.* [60] extended GP to compute joint likelihood of multiple loci under null and alternative hypotheses. Recently, Schraiber *et al.* [54] proposed a Bayesian framework to estimate parameters using Monte Carlo Markov chain sampling.

While existing methods have been successfully applied to their corresponding application, they make some assumptions which may not hold in E&R studies. First, they assume that the underlying population size is large, so it is reasonable to model dynamics of allele frequencies using continuous state models. A number of existing methods were originally designed to process wide time spans such as ancient DNA studies. Finally, they assume that input data is in the form of unbiased allele frequencies, which may not be valid for shotgun sequencing experiments.

Here, we consider a Hidden Markov Model (HMM), similar to Williamson *et al.* [67] and Bollback et *al.*’s [11] but under a “small-population-size” regime. Specifically, we use a discrete state (frequency) model. We show that for small population sizes, discrete models can compute likelihood exactly, which improves statistical performance, especially for short time-span experiments. Additionally, we add another level of sampling-noise to the traditional HMM model, allowing for heterogeneous ascertainment bias due to uneven coverage among variants. We show that for a wide range of parameters, Clear provides higher power for detecting selection, estimates model parameters consistently, and localizes favored allele more accurately compared to the state-of-the-art methods, while being computationally efficient.

## 2 Materials and Methods

Consider a panmictic diploid population with fixed size of *N* individuals. Let *ν* = {*ν*_*t*_}_*t*∈ 𝒯_ be frequencies of the derived allele at generations *t* ∈ 𝒯 for a given variant, where at generations 𝒯 = {τ_*i*_ : 0 ≤ τ_0_ < τ_1_,… < τ_*T*_} samples of *n* individuals are chosen for pooled sequencing. The experiment is replicated *R* times. We denote allele frequencies of the *R* replicates by the set {*ν*}_*R*_. To identify the genes and variants that are responding to selection pressure, we use the following procedure:

(i) **Estimating population size.** The procedure starts by estimating the effective population size, *N***̂**, under the assumption that much of the genome is evolving neutrally.
(ii) **Estimating selection parameters.** For each polymorphic site, selection and dominance parameters *s*, *h* are estimated so as to maximize the likelihood of the time series data, given *N***̂**
(iii) **Computing likelihood statistics.** For each variant, a log-odds ratio of the likelihood of selection model (*s* > 0) to the likelihood of neutral evolution/drift model is computed. Likelihood ratios in a genomic region are combined to compute the Clear statistic for the region.
(iv) **Hypothesis testing.** An empirical null distribution of the Clear statistic is calculated using genome-wide drift simulations, and used to compute *p*-values and thresholds for a specified FDR. We perform single locus hypothesis testing within selected regions to identify significant variants and report genes that intersect with the selected variants.

These steps are described in detail below.

### 2.1 Estimating Population Size

Methods for estimating population sizes from temporal neutral evolution data have been developed [2, 11, 33, 60, 67]. Here, we aim to extend these models to explicitly model the sampling noise that arise in pool-seq data. Specifically, we model the variation in sequence coverage over different locations, and the noise due to sequencing only a subset of the individuals in the population. In addition, many existing methods [11, 23, 60, 62] are designed for large populations, and model frequency as a continuous quantity. We show that smooth approximations may be inadequate for small populations, low starting frequencies and sparse sampling (in time) that are typical in experimental evolution (see Results, Fig 3A-C, and Fig 2). To this end, we model the Wright-Fisher Markov process for generating pool-seq data (Fig S1) via a discrete HMM (Fig 1-B). We start by computing a likelihood function for the population size given neutral pool-seq data.

**Fig 1:**
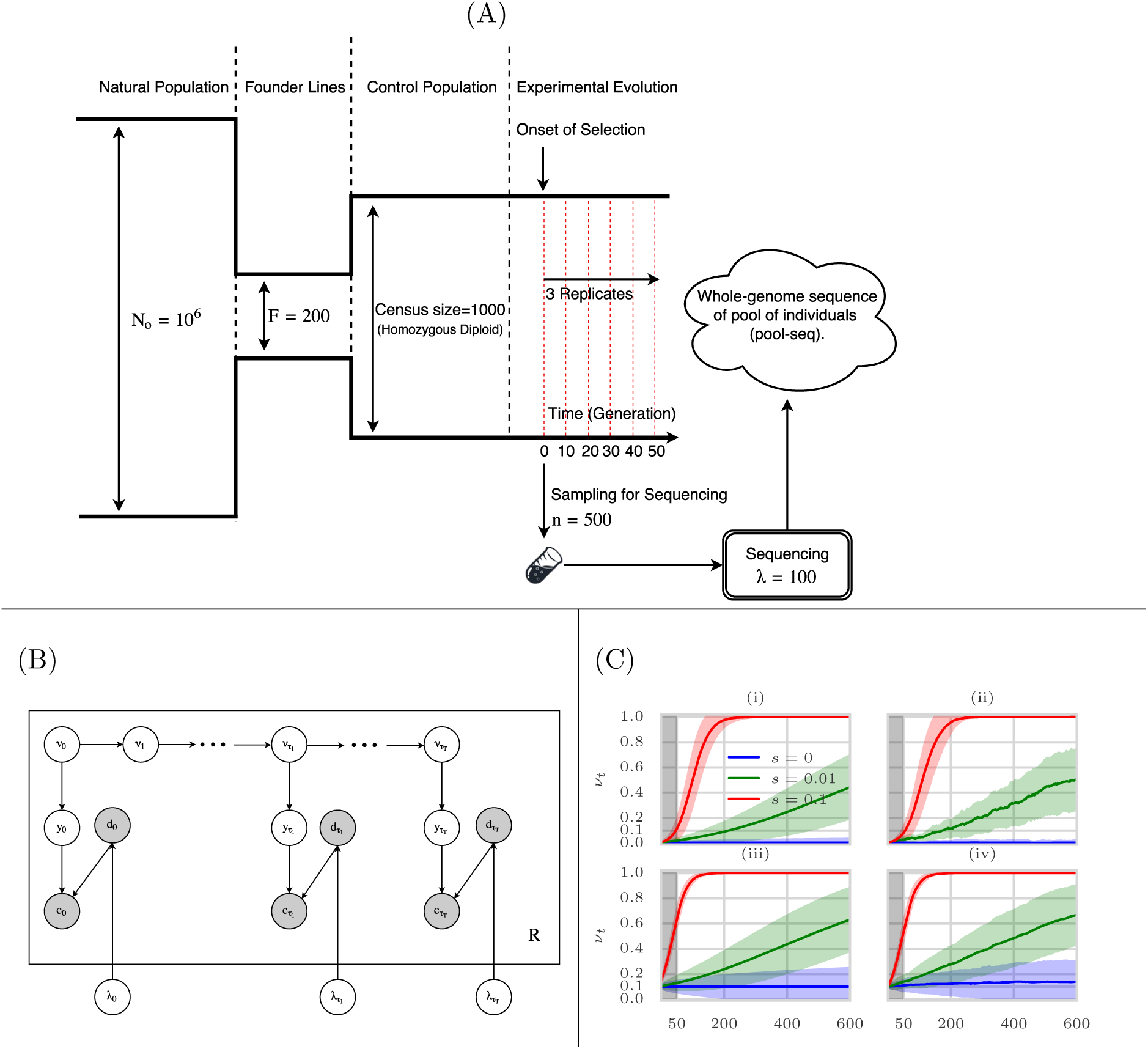
Evolve and Resequence Selection Experiments on *D. melanogaster*. (A) Typical configuration in which time-series data is collected for *D. melanogaster.* A small set of founder lines (*F* = 200) is selected from a large population (*N_o_* = 10^6^), and used to create a sub-population of isofemale lines. Multiple replicates of the population are evolved and resequenced to collect time-series genomic data. For sequencing, *n* individuals are randomly sampled and sequenced with coverage *λ*. (B) Graphical model showing dependence of the random variables in the singlelocus model used to compute Clear statistics. Observed variables, c (derived allele read count) and d (total read count) are shaded. The variables *v, y,* λ denote allele frequency, sampled allele frequency, and mean sequencing coverage, respectively. (C) Mean and 95% confidence interval of the theoretical (i,iii) and empirical (ii,iv) trajectories of the favored allele for hard (i,ii) and soft (iii,iv) sweep scenarios and *N* = 1000. The first 50 generations are shaded in gray to represent the sampling span of sampling in short-term experiments, illustrating the difficulty in predicting selection at early stages of selective sweep.

**Fig 2:**
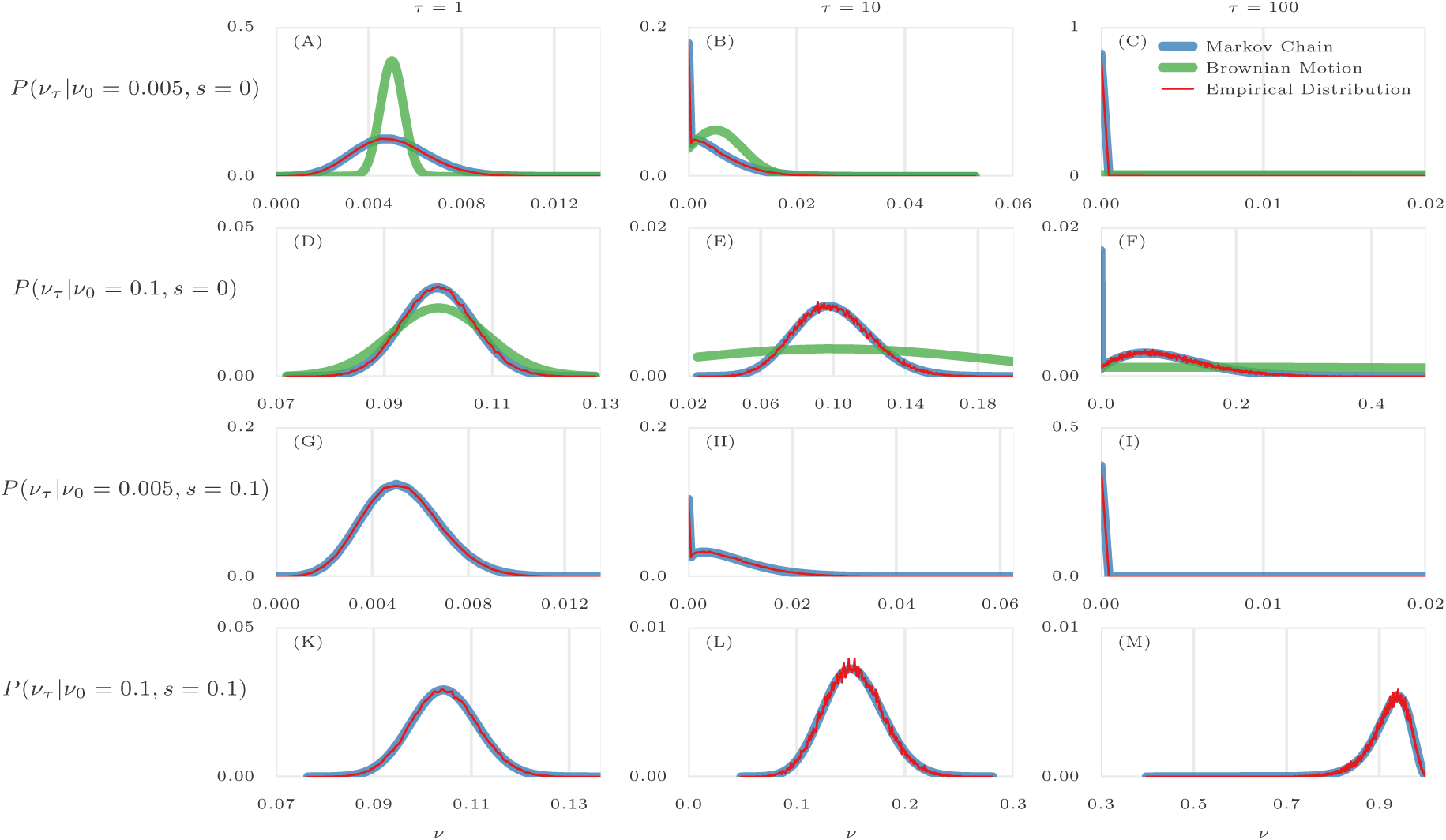
Comparison of empirical distributions of allele frequencies (red) versus predictions from Brownian Motion (green), and Markov chain (blue). Comparison of empirical and theoretical distributions under neutral evolution (panels A-F) and selection (panels G-M) with different starting frequencies *ν*_*0*_ ∈ {0.005,0.1} and sampling times of 𝒯 = {0, τ}, where τ ∈ {1,10,100}. For each panel, the empirical distribution was computed over 100,000 simulations. Brownian motion (Gaussian approximation) provides poor approximations when initial frequency is far from 0.5 (A) or sampling is sparse (B,C,E,F). In addition, Brownian motion can only provide approximations under neutral evolution. In contrast, Markov chain consistently provides a good approximation in all cases.

**Fig 3:**
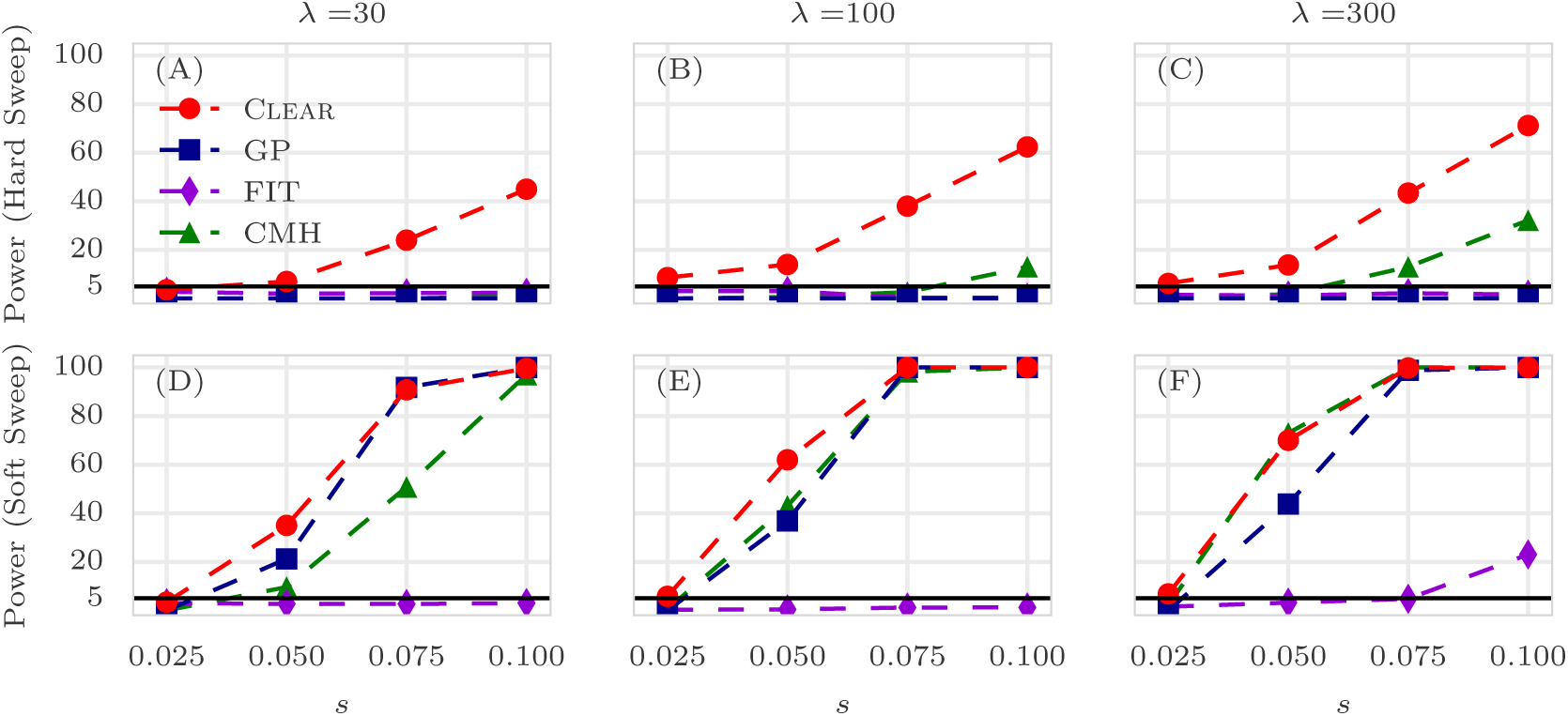
Power calculations for detection of selection. Detection power for Clear(𝓗), Frequency Increment Test (FIT), Gaussian Process (GP), and CMH under hard (A-C) and soft sweep (D-F) scenarios. *λ*, *s* denote the mean coverage and selection coefficient, respectively. The *y*-axis measures power – sensitivity with false positive rate FPR ≤ 0.05 – for 2, 000 simulations with *N* = 1, 000, *L* = 50Kbp. The horizontal line reflects the power of a random classifier. In all simulations, 3 replicates are evolved and sampled at generations 𝒯 = {0,10, 20, 30, 40, 50}.

#### Likelihood for Neutral Model

We model the allele frequency counts 2*Nν*_*t*_ as being sampled from a Binomial distribution. Specifically,

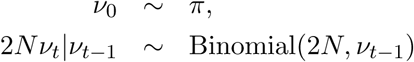

where π is the global distribution of allele frequencies in the base population. Here we simply assume is π is the site frequency spectrum of fixed sized neutral population Fig S2. Note that π may depend on the demographic history of the founder lines.

To estimate frequency after τ transitions, it is enough to specify the 2*N* × 2*N* transition matrix *P*^(τ)^, where *P*^(τ)^[*i*,*j*] denotes probability of change in allele frequency from *i*/*2N* to *j*/*2N* in τ generations:

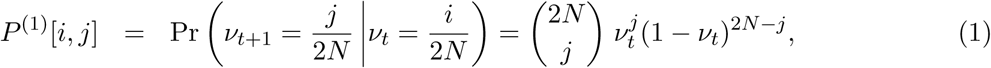

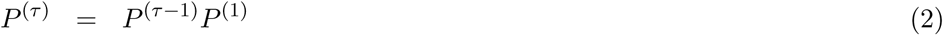

Furthermore, in an E&R experiment, *n* ≤ *N* individuals are randomly selected for sequencing. The sampled allele frequencies, {*y_t_*}_*t*∈𝒯_, are also Binomially distributed

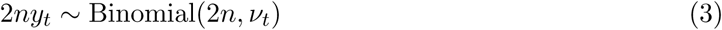

We introduce the 2*N* × 2*n* sampling matrix Y, where *Y*[*i*, *j*] stores the probability that the sample allele frequency is *j*/2*n* given that the true allele frequency is *i*/2*N*.

We denote the pool-seq data for that variant as {*x_t_* = <*c_t_*,*d_t_*>}_*t*∈ 𝒯_ where *d_t_,c_t_* represent the coverage, and the read count of the derived allele, respectively. Let {*λ*_*t*_}_*t* ∈ 𝒯_ be the sequencing coverage at different generations. Then, the observed data are sampled according to

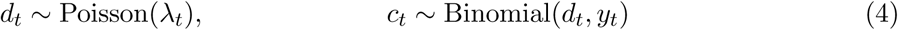

The emission probability for a observed tuple *x_t_* = <*d_t_, c_t_*> is

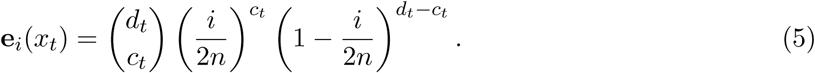

For 1 ≤ *t* ≤ *T,* 1 ≤ j ≤ 2*N*, let *α_t,j_* denote the probability of emitting *x*_1_*,x*_2_,…,*x_t_* and reaching state *j* at τ_*t*_. Then, *α*_*t*_ can be computed using the forward-procedure [19]:

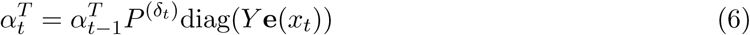

where δ_*t*_ = τ_*t*_ − τ_*t*-1_. The joint likelihood of the observed data from *R* independent observations is given by

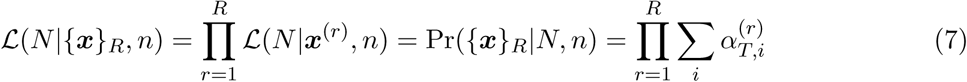

where ***x*** = {*x_t_*}_*t*∈𝒯_. The graphical model and the generative process for which data is being generated is depicted in Fig 1-B and Fig S1, respectively.

Finally, the last step is to compute an estimate *N̂* that maximizes the likelihood of all *M* variants in whole genome. Let 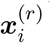 denote the time-series data of the *i*-th variant in replicate *r.* Then,

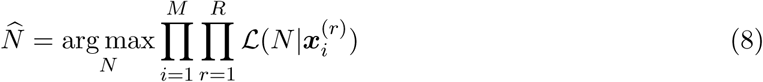

### 2.2 Estimating Selection Parameters

#### Likelihood for Selection Model

Assume that the site is evolving under selection constraints *s* ∈ ℝ, *h* ∈ ℝ_+_, where s and *h* denote selection strength and dominance parameters, respectively. By definition, the relative fitness values of genotypes 0|0, 0|1 and 1|1 are given by *w*_00_ = 1, *w*_01_ = 1 + *hs* and *w*_11_ = 1 + *s*. Then, *ν*_*t*+_, the frequency at time τ _*t*_ + 1 (one generation ahead), can be estimated using:

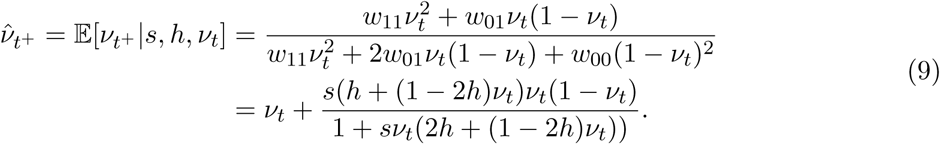

The machinery for computing likelihood of the selection parameters is identical to that of population size, except for transition matrices. Hence, here we only describe the definition transition matrix *Q_s_,_h_* of the selection model. Let 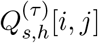 denote the probability of transition from *i*/2*N* to *j/*2*N* in τ generations, then (See [20], Pg. 24, Eqn. 1.58-1.59):

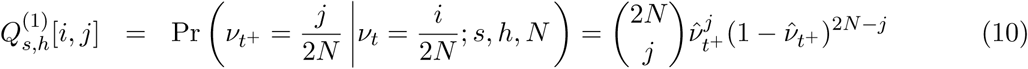

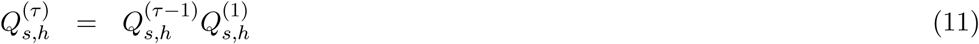

The maximum likelihood estimates are given by

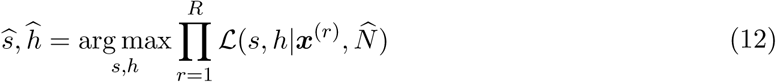

Using grid search, we first estimate *N* (Eq. 8), and subsequently, we estimate parameters *s*, *h* (Eq. 12, Fig S3). By broadcasting and vectorizing the grid search operations across all variants, the genome scan on millions of polymorphisms can be done in significantly smaller time than iterating a numerical optimization routine for each variant(see Results and Fig 4).

**Fig 4:**
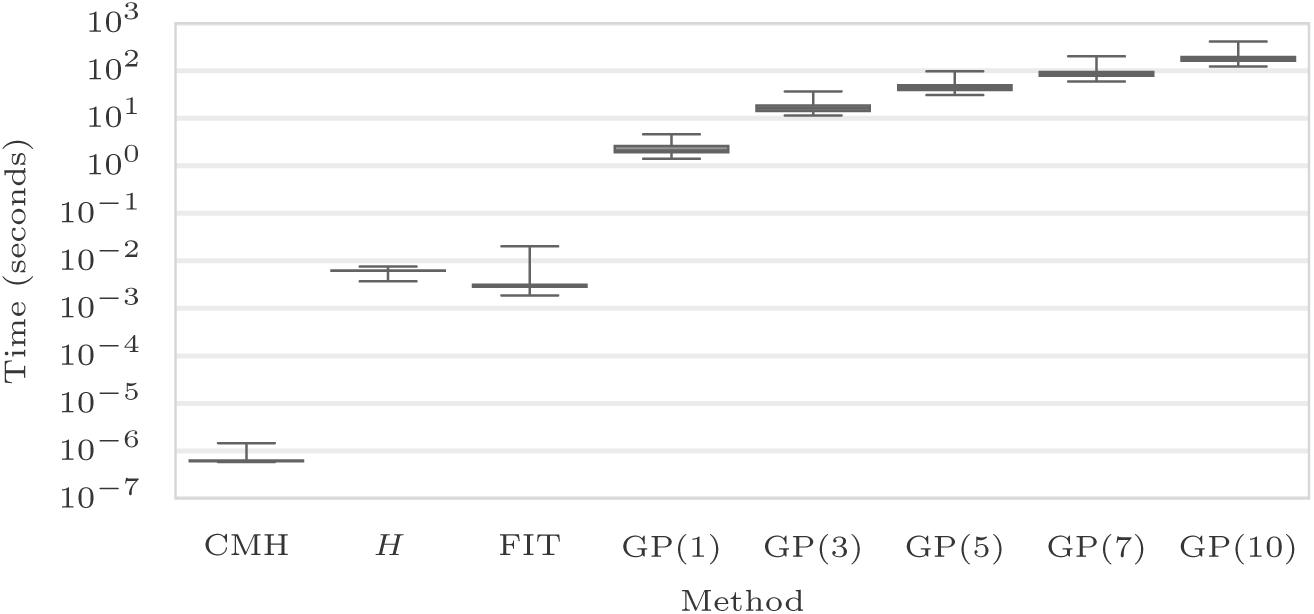
Running time. Box plots of running time per variant (CPU-secs.) of Clear(𝓗), CMH, FIT, and GP with single, 3, 5, 7, and 10 loci over 1000 simulations conducted on a workstation with Intel Core i7 processor. The average running time for each method is shown on the x-axis. In all simulations, 3 replicates are evolved and sampled at generations 𝒯 = {0; 10; 20; 30; 40; 50}.

### 2.3 Empirical Likelihood Ratio Statistics

The likelihood ratio statistic for testing directional selection, to be computed for each variant, is given by

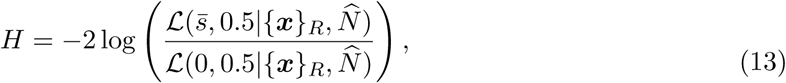

where *s̅* =arg max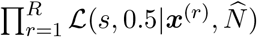. Similarly we can define a test statistic for testing if selection is dominant by

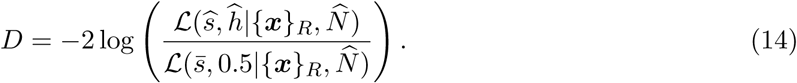

While extending the single-locus WF model to a multiple linked-loci can improve the power of the model [60], it is computationally and statistically expensive to compute exact likelihood. In addition, computing linked-loci joint likelihood requires haplotype resolved data, which pool-seq does not provide. Here, similar to Nielsen *et al* [45], we calculate *composite likelihood ratio* score for a genomic region.

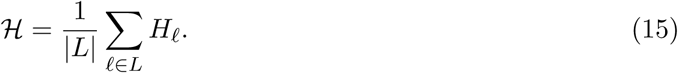

where *L* is a collection of segregating sites and *H*_ℓ_ is the likelihood ratio score based for each variant ℓ in *L*. The optimal value of the hyper-parameter *L* depends upon a number of factors, including initial frequency of the favored allele, recombination rates, linkage of the favored allele to neighboring variants, population size, coverage, and time since the onset of selection (duration of the experiment). In S1 Text, we provide a heuristic to compute a reasonable value of *L*, based on experimental data.

We work with a normalized value of 𝓗, given by

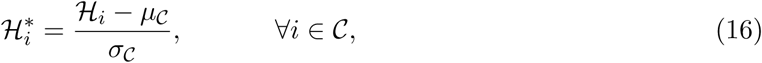

where *μ_C_* and *σ_C_* are the mean and standard deviation of 𝓗 values in a large region 𝒞. We found different chromosomes to have different distribution of 𝓗_*i*_ values, and therefore decided to use single chromosomes as 𝒞.

### 2.4 Hypothesis Testing

#### Single-Locus tests

Under neutrality, Log-likelihood ratios can be approximated by *𝒳*^2^ distribution [66], and *p*-values can be computed directly. However, Feder *et al.* [23] showed that when the number of independent samples (replicates) is small, *𝒳*^2^ is a crude approximation to the true null distribution and results in more false positive. Following their suggestion, we first compute the empirical null distribution using simulations with the estimated population size (See Fig S1). The empirical null distribution of statistic *H* is used to compute *p*-values as the fraction of null values that exceed the test score. Finally, we use Storey and Tibshirani’s method [59] to control for False Discovery Rate in multiple testing.

#### Composite likelihood tests

Similar to single-locus tests, we compute the null distribution of the 𝓗* statistic using whole-genome simulations with the estimated population size, and subsequently compute FDR. The simulations for generating the null distribution of 𝓗* are described next.

### 2.5 Simulations

We use the same simulation procedure for two purposes. First, we use them to test the power of Clear against other methods in small genomic windows. Second, we use the simulations to generate the distribution of null values for the statistic to compute empirical *p*-values. We mainly chose parameters that are relevant to *D. melanogaster* experimental evolution [35]. See also Fig 1-A for illustration.

I. **Creating initial founder line haplotypes.** Using `msms` [21], we created neutral populations for *F* founding haplotypes with command $./msms <F> 1 -t <2*μ*WNe> -r <2rNeW> <W>, where *F* = 200 is number of founder lines, *N_o_* = 10^6^ is effective founder population size, *r* = 2 × 10^−8^ is recombination rate, *μ* = 2 × 10^−9^ is mutation rate. The window size *W* is used to compute θ = 2 *μ N_o_W* and ρ = 2*N_o_rW*. We chose *W* = 50Kbp for simulating individual windows for performance evaluations, and *W* = 20Mbp for simulating *D*. melanogaster chromosomes for *p*-value computations.
II **Creating initial diploid population.** An initial set of *F =* 200 haplotypes was created from step I, and duplicated to create *F* homozygous diploid individuals to simulate generation of inbred lines. *N* diploid individuals were generated by sampling with replacement from the *F* individuals.
III **Forward Simulation.** We used forward simulations for evolving populations under selection. We also consider selection regimes which the favored allele is chosen from standing variation (not *de novo* mutations). Given initial diploid population, position of the site under selection, selection strength s, number of replicates *R* = 3, recombination rate *r* = 2 × 10^−8^ and sampling times 𝒯 = {0,10, 20, 30, 40, 50}, simuPop [48] was used to perform forward simulation and compute allele frequencies for all of the *R* replicates. For hard sweep (respectively, soft sweep) simulations we randomly chose a site with initial frequency of *ν*_0_ = 0.005 (respectively, *ν*_0_ = 0.1) to be the favored allele. For generating the null distribution with drift for *p*-value computations, we used this procedure with *s* = 0.
IV. **Sequencing Simulation.** Given allele frequency trajectories we sampled depth of each site in each replicate identically and independently from Poisson(*λ*), where *λ* ∈ {30,100,300} is the coverage for the experiment. Once depth *d* is drawn for the site with frequency *ν*, the number of reads c carrying the derived allele are sampled according to Binomial(*d*,*ν*). For experiments with finite depth the tuple <*c*, *d*> is the input data for each site.

## 3 Results

### Modeling Allele Frequency Trajectories in Small Populations

We first tested the goodness of fit of the discrete versus continuous models in modeling allele frequency trajectories, under general E&R parameters. For this purpose, we conducted 100K simulations with two time samples 𝒯 = {0, τ} where τ ∈ {1,10,100} is the parameter controlling the density of sampling in time. In addition, we repeated simulations for different values of starting frequency *ν*_0_ ∈ {0.005,0.1} (i.e., hard and soft sweep) and selection strength *s* ∈ {0,0.1} (i.e., neutral and selection). Then, given initial frequency *ν*_0_, we computed the expected distribution of the frequency of the next sample *ν* _τ_ under two models to make a comparison. Fig 2A-F shows that Brownian motion (continuous model) is inadequate when *ν*_0_ is far from 0.5, or when sampling times are sparse (τ > 1). If the favored allele arises from standing variation in a neutral population, it is unlikely to have frequency close to 0.5, and the starting frequencies are usually much smaller (see Fig S2). Moreover, in typical *D. melanogaster* experiments for example, sampling is sparse. Often, the experiment is designed so that 10 ≤ τ ≤ 100 [26, 35, 46, 71].

In contrast to the Brownian motion approximation, discrete Markov chain predictions (Eq. 11) are highly consistent with empirical data for a wide range of simulation parameters (Fig 2A-M). Moreover, the discrete markov chain can be modified to model the case when the the allele is under selection.

### Detection Power

We compared the performance of Clear against other methods for detecting selection. For each method we calculated detection power as the percentage of true-positives identified with false-positive rate ≤ 0.05. For each configuration (specified with values for selection coefficient *s*, starting allele frequency *ν*_0_ and coverage *λ*), power of each method is evaluated over 2000 distinct simulations, half of which modeled neutral evolution and the rest modeled positive selection.

We compared the power of Clear with Gaussian process (GP) [60], FIT [23], and CMH [1] statistics. FIT and GP convert read counts to allele frequencies prior to computing the test statistic. Clear shows the highest power in all cases and the power stays relatively high even for low coverage (Fig 3 and Table S1). In particular, the difference in performance of Clear with other methods is pronounced when starting frequency is low. The advantage of Clear stems from the fact that favored allele with low starting frequency might be missed by low coverage sequencing. In this case, incorporating the signal from linked sites becomes increasingly important. We note that methods using only two time points, such as CMH, do relatively well for high selection values and high coverage. However, the use of time-series data can increase detection power in low coverage experiments or when starting frequency is low. Moreover, time-series data provide means for estimating selection parameters *s, h* (see below). Finally, as Clear is robust to change of coverage, our results (Fig 3B,C) suggest that taking many samples with lower coverage is preferable to sparse sampling with higher coverage.

### Site-identification

In general, localizing the favored variant, using pool-seq data is a nontrivial task due to extensive linkage disequilibrium [61]. To measure performance, we sorted variants by their H scores and computed rank of the favored allele for each method. For each setting of ν_0_ and s, we conducted 1000 simulations and computed the rank of the favored mutation in each simulation. The cumulative distribution of the rank of the favored allele in 1000 simulation for each setting (Fig 5) shows that Clear outperforms other statistics.

**Fig 5:**
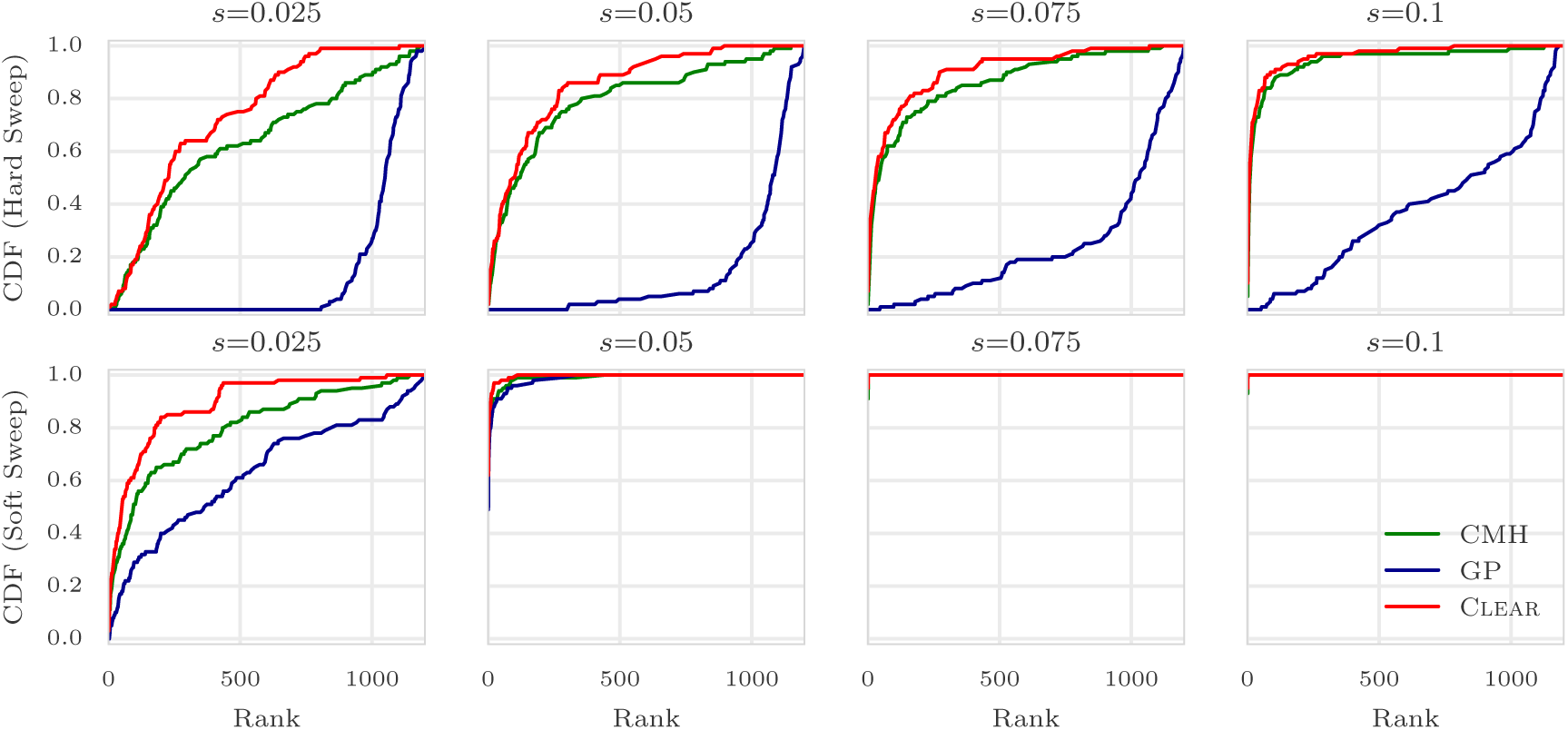
Ranking performance for 100× coverage. Cumulative Distribution Function (CDF) of the distribution of the rank of the favored allele in 1000 simulations for Clear (*H*), Gaussian Process (GP), CMH, and Frequency Increment Test (FIT), for different values of selection coefficient s and initial carrier frequency. Note that the individual variant Clear score (*H*) is used to rank variants. The Area Under Curve (AUC) is computed as an overall quantitative measure to compare the performance of methods for each configuration. In all simulations, 3 replicates are evolved and sampled at generations 𝒯 = {0,10,20,30,40, 50}.

An interesting observation is revisiting the contrast between site-identification and detection [39, 61]. When selection strength is high, detection is easier (Fig 3A-F), but site-identification is harder, due to the high LD between flanking variants and the favored allele (Fig 5A-F). Moreover, site-identification becomes more difficult whenever the initial frequency of the favored allele is low, i.e., at the onset of selection, LD between favored allele and its nearby variants is high. For example, when coverage *λ* = 100 and selection coefficient *s* = 0.1, the detection power is 75% for hard sweep, but 100% for soft sweep (Fig 3B-E). In contrast, the favored site was ranked as the top in 14% of hard sweep cases, compared to and 95% of soft sweep simulations.

### Estimating Parameters

Clear estimates effective population size *N̂* and selection parameters, *ŝ* and *ĥ*, as a byproduct of the hypothesis testing. We computed bias of selection fitness (*s* – *ŝ*) and dominance (*h* – *ĥ*) for of Clear and GP for 1000 simulations in each setting. The distribution of the error (bias) for 100 × coverage is presented in Fig 6 for different configurations. Fig S4 and Fig S5 provide the distribution of estimation errors for 30 ×, and 300 × coverage, respectively. For hard sweep, Clear provides estimates of s with lower variance of bias (Fig 6A). In soft sweep, GP and Clear both provide unbiased estimates of *s* with low variance (Fig 6B). Fig 6C-D shows that Clear provides unbiased estimates of *h* as well when *h* ∈ {0,0.5,1, 2} and *s* = 0.1. We also tested if Clear provide unbiased estimates of *N*, by estimating population size on 1000 simulations when N ∈ {200, 600,1000}. As shown in Fig 7A-C, maximum likelihood is attained at true value of the parameter.

**Fig 6:**
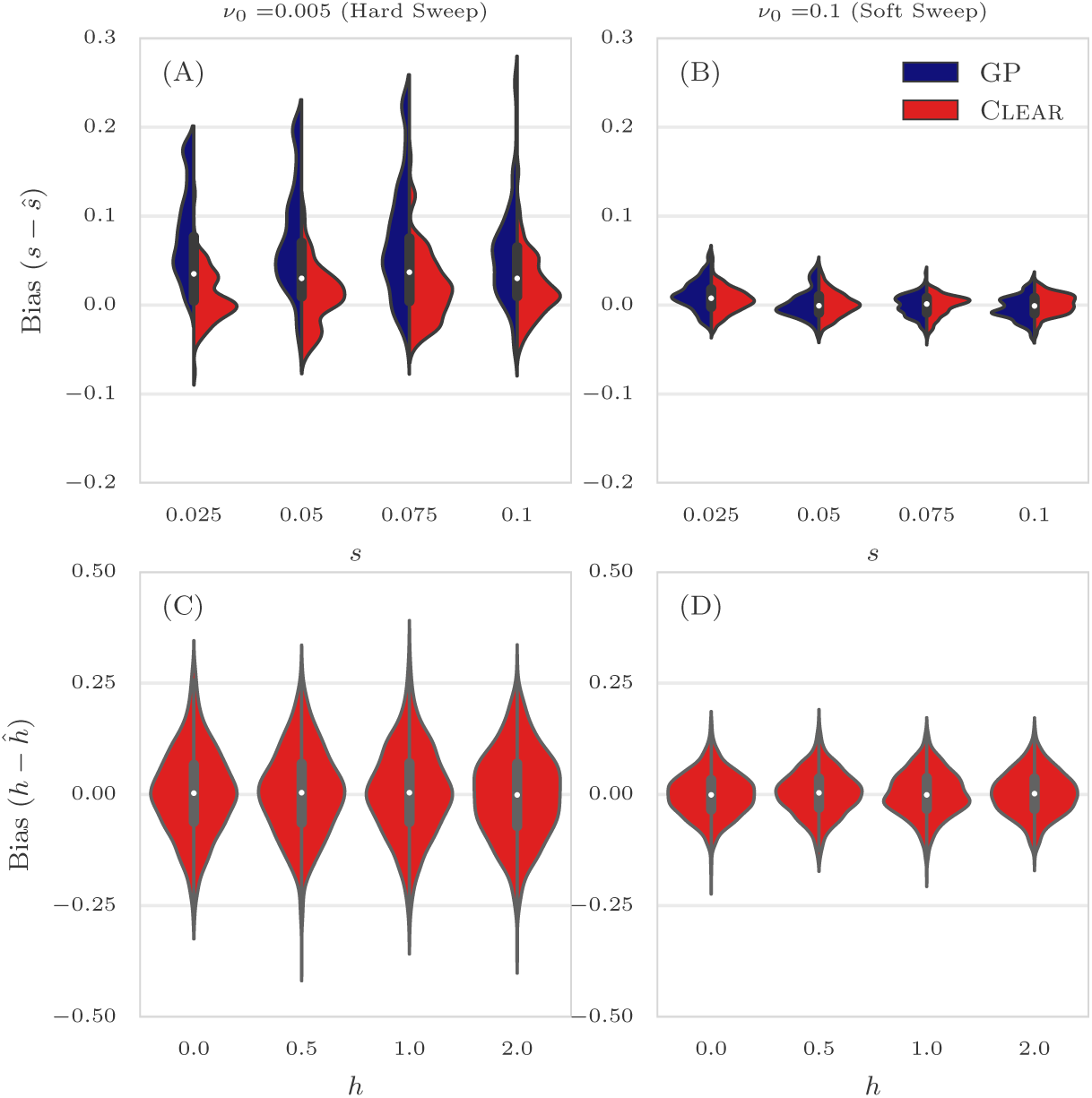
Distribution of bias for 100 × coverage. The distribution of bias (*s* – *ŝ*) in estimating selection coefficient over 1000 simulations using Gaussian Process (GP) and Clear (*H*) is shown for a range of choices for the selection coefficient *s* and starting carrier frequency *ν*_0_when coverage *λ* = 100 (Panels A,B). GP and Clear have similar variance in estimates of s for soft sweep, while Clear provides lower variance in hard sweep. Also see Table S2. Panels C,D show the variance in the estimation of *h*. In all simulations, 3 replicates are evolved and sampled at generations 𝒯 = {0,10,20,30, 40, 50}.

**Fig 7:**
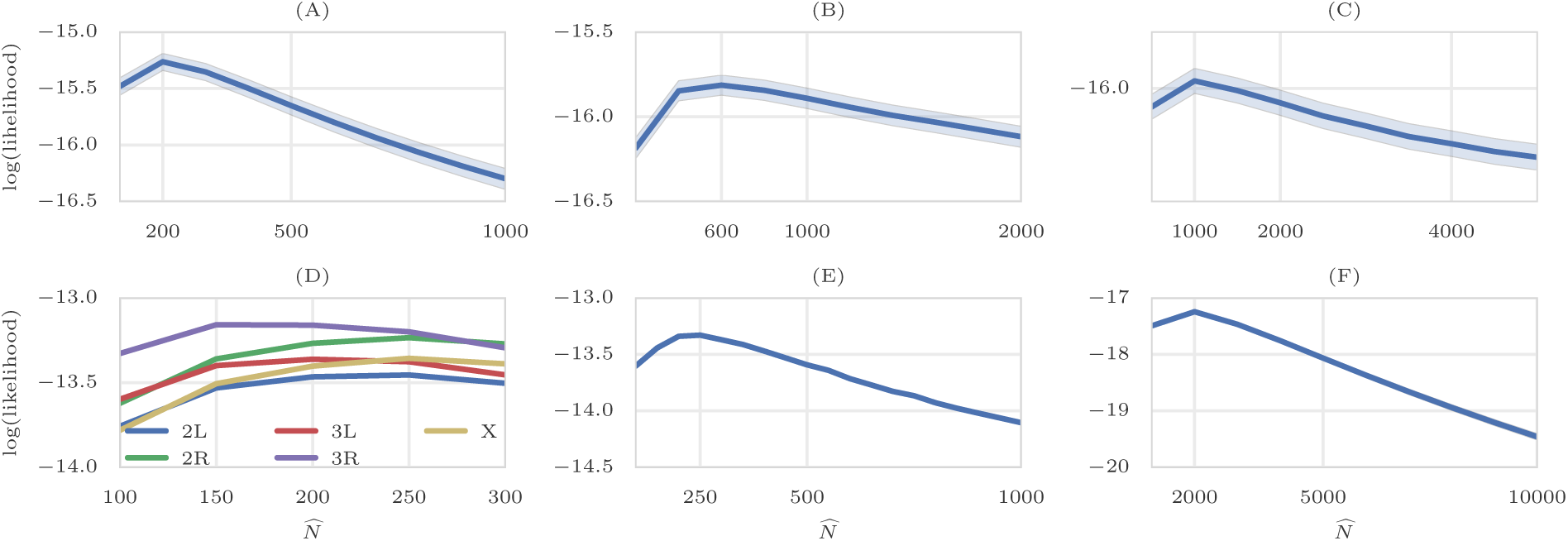
Maximum likelihood Estimates of *N*. Mean and 95% confidence interval of likelihoods of *N* on simulated data with *N* = 200 (A), *N* = 600(B), and *N* = 1000 individuals, over 1000 simulations. Chromosome-wise (D) and genome-wide (E) likelihood of population size for data from a study of *D. melanogaster* adaptation to alternating temperatures. Likelihood of the Chromosome 3R is attained at 150, while genome-wide maximum likelihood estimate for population size is 250. (F) Likelihood of the population size with respect to all the variants in the yeast dataset. Despite large census population size (10^6^ – 10^7^ [14]), this dataset exhibits much smaller effective population size (*N̂* = 2000).

### Running Time

As Clear does not compute exact likelihood of a region (i.e., does not explicitly model linkage between sites), the complexity of scanning a genome is linear in number of polymorphisms. Calculating score of each variant requires and 𝒪(*TRN*^3^) computation for 𝓗. However, most of the operations are can be vectorized for all replicates to make the effective running time for each variant. We conducted 1000 simulations and measured running times for computing site statistics *H*, FIT, CMH and GP with different number of linked-loci. Our analysis reveals (Fig 4) that Clear is orders of magnitude faster than GP, and comparable to FIT. While slower than CMH on the time per variant, the actual running times are comparable after vectorization and broadcasting over variants (see below).

These times can have a practical consequence. For instance, to run GP in the single locus mode on the entire pool-seq data of the *D. melanogaster* genome from a small sample (≈1.6M variant sites), it would take 1444 CPU-hours (≈ 1 CPU-month). In contrast, after vectorizing and broadcasting operations for all variants operations using numba package, Clear took 75 minutes to perform an scan, including precomputation, while the fastest method, CMH, took 17 minutes.

#### 3.1 Analysis of a *D. melanogaster* Adaptation to Alternating Temperatures

We applied Clear to the data from a study of *D*. *melanogaster* adaptation to alternating temperatures [26, 46], where 3 replicate samples were chosen from a population of *D. melanogaster* for 59 generations under alternating 12-hour cycles of hot stressful (28°C) and non-stressful (18°C) temperatures and sequenced. In this dataset, sequencing coverage is different across replicates and generations (see S2 Fig of [60]) which makes variant depths highly heterogeneous (Fig S8).

We first filtered out heterochromatic, centromeric and telomeric regions [25], and those variants that have collective coverage of more that 1500 in all 13 populations: three replicates at the base population, two replicates at generation 15, one replicate at generation 23, one replicate at generation 27, three replicates at generation 37 and three replicates at generation 59. After filtering, we ended up with 1,605,714 variants.

Next, we estimated genome-wide population size N̂ = 250 (Fig 7-E) which is consistent with previous studies [33, 46]. The likelihood curves of Clear are sharper around the optimum compared to that of Bollback et. al [11]’s method (see Supplementary Fig. 1 in [46]). Also, chromosomes 3L and 3R appear to have smaller population size Fig 7-D, *N̂* = 200,150, respectively. Others have made similar observations on this data. In particular, Jonas *et al.* [33] shown that the chromosome-wise population size varies even more when it is computed for each replicate separately (see Table 1 in [33]). For instance, N̂ is 131 for chromosome 3R replicate 1, while it is 328 for chromosome X replicate 2.

While it would be ideal to compute Clear statistic for each replicate and chromosome separately, computing empirical *p*-values and significant regions become computationally intensive as empirical null distribution of each replicate and each chromosome needs to be computed. Hence, we use a single genome-wide estimate *N̂* = 250 in all analyses, but we normalize statistic 𝓗* separately for each chromosome.

We use a heuristic calculation (See S1 Text) to choose the sliding window size *L* as the distance where the LD between the favored mutation and a site *L*/2bp away remains strong. For *D. melanogaster* parameters, we obtained *L* = 30kbp. We computed the normalized test statistic 𝓗* on sliding windows of size of 30Kbp and step size of 5Kbp over the genome (See Fig 8-A).

**Fig 8:**
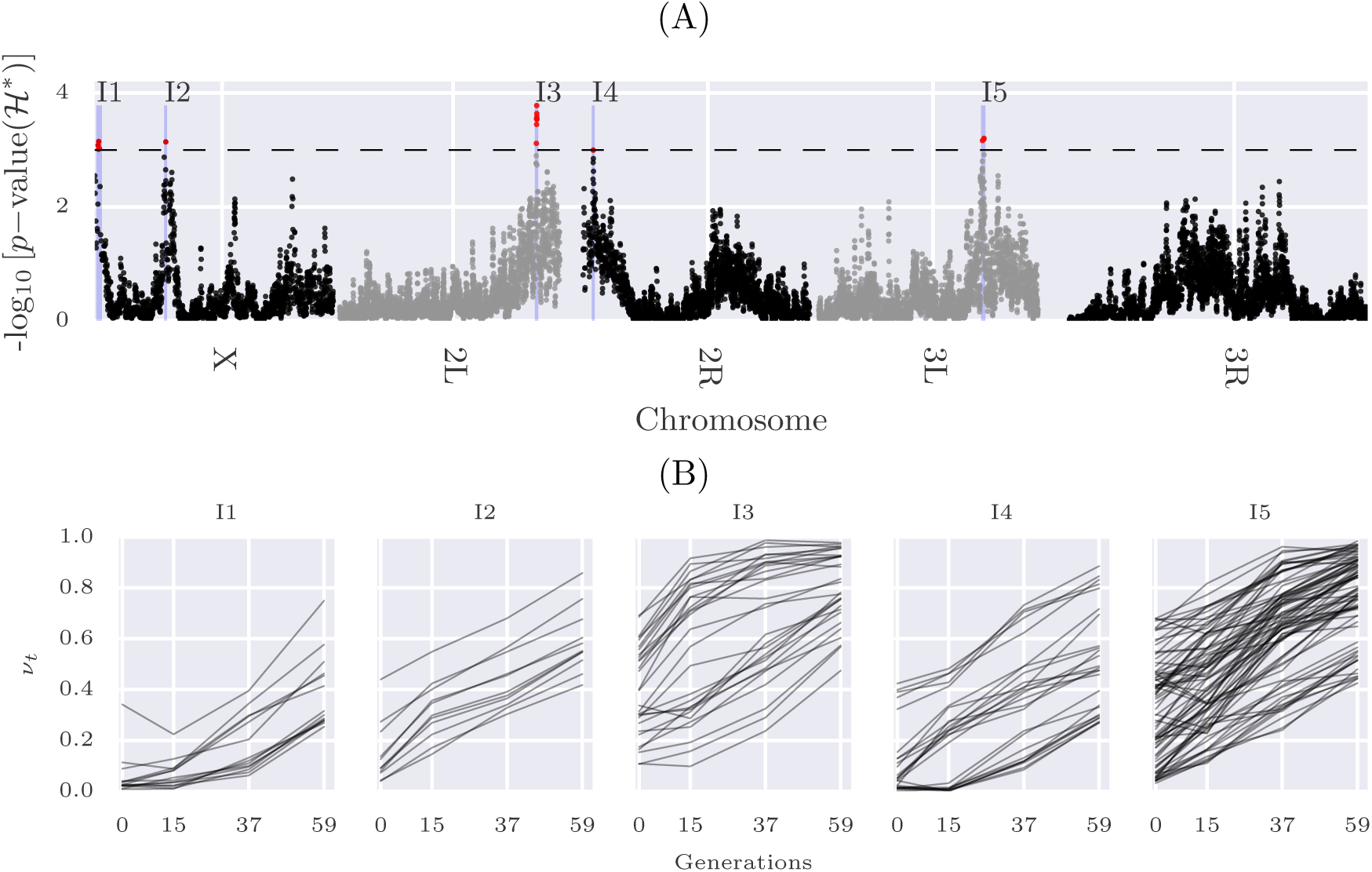
Scan of Clear statistic on data from a study of *D. melanogaster* adaptation to alternating temperatures. (A) Manhattan plot of scan for 𝓗*\** statistic over the genome. The dashed line represents cutoff for genome-wide FDR≤ 0.05, and identifies 5 contiguous intervals, I1-I5, which are shaded in blue. (B) Trajectories of the selected variants within intervals I1-I5.

**Fig 9:**
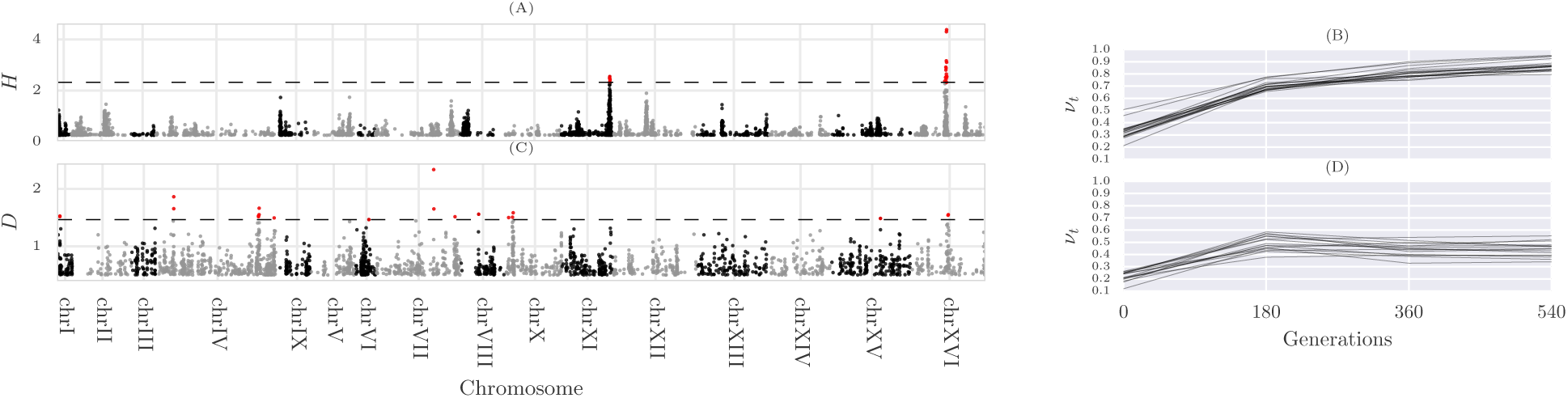
Single locus analysis of the yeast outcrossing populations. Manhattan plot of scan for testing directional selection (A) and dominant selection (C). The dashed line represents cutoff for genome-wide FDR≤ 0.05. Trajectories of the selected variants are depicted in panels (B) and (D).

Empirical null distribution of 𝓗* was estimated by creating 100 whole genome simulations (400K statistic values) as described in Section 2.5. Then, *p*-value of the test statistic in each region in the experimental data was calculated as the fraction of the null statistic values that are greater than or equal to the test statistic(see Fig S9). After correcting for multiple testing, we identified 5 contiguous intervals (Fig 8) satisfying FDR≤ 0.05, and covering 2, 829 polymorphic sites. We further performed single-locus hypothesis testing on the 2,829 sites to identify 174 individual variants with FDR ≤ 0.01 (Fig 8-B).

The final set of 174 variants fall within 32 genes(Table S3) including many Serine inhibitory proteases (serpins), and other genes involved in endocytosis. Recycling of synaptic vesicles is seen to be blocked at high temperature in temperature sensitive Drosophila mutants [36]. This is also supported by GO enrichment analysis, where a single GO term ‘inhibition of proteolysis’ is found to enriched (corrected *p*-value:0.0041). To test for dominant selection, we computed D statistic on simulated neutral and experimental data, and computed *p*-values accordingly. After correcting for multiple testing, 96 variants were discovered with FDR ≤ 0.01 (Fig S10).

#### 3.2 Analysis of Outcrossing Yeast Populations

We also applied Clear to 12 replicate samples of outcrossing yeast populations [14], where samples are taken at generations 𝒯 = {0,180,360, 540}. We observed a significant variation in the genome-wide site frequency spectrum of certain populations over different time points for some replicates (Fig S11). The variation does not have an easily identifiable cause. Therefore, we focused analysis on seven replicates *r* ∈ {3, 7,8,9,10,11,12} with genome-wide site-frequency spectrum over the time range (Fig S12).

We estimated population size to be *N̂* = 2000 haplotypes, and computed *ŝ*, *ĥ* and *H* statistic accordingly. To compute *p*-values, we created 1M single-locus neutral simulations according to experimental data’s initial frequency and coverage. By setting FDR cutoff to 0.05, only 18 and 16 variants show significant signal for directional and dominant selection, respectively (Fig S10). Selected variants for directional selection are clustered in two regions, which match 2 of the 5 regions (regions C and E in Fig. 2-a in [14]) identified by Burke *et al.* in their preliminary analysis.

## 4 Discussion

We developed a computational tool, Clear, that can detect regions and variants under selection E&R experiments. Using extensive simulations, we show that Clear outperforms existing methods in detecting selection, locating the favored allele, and estimating model parameters. Also, while being computationally efficient, Clear provide means for estimating populations size and hypothesis testing.

Many factors such as small population size, finite coverage, linkage disequilibrium, finite sampling for sequencing, duration of the experiment and the small number of replicates can limit the power of tools for analyzing E&R. Here, by an discrete modeling, Clear estimates population size, and provides unbiased estimates of *s*, *h*. It adjusts for heterogeneous coverage of pool-seq data, and exploits presence of linkage within a region to compute composite likelihood ratio statistic.

It should be noted that, even though we described Clear for small fixed-size populations, the statistic can be adjusted for other scenarios, including changing population sizes when the demography is known. For large populations, transitions can be computed on sparse data structures, as for large *N* the transition matrices become increasingly sparse. Alternatively, frequencies can be binned to reduce dimensionality.

The comparison of hard and soft sweep scenarios showed that initial frequency of the favored allele can have an nontrivial effect on the statistical power for identifying selection. Interestingly, while it is easier to detect a region undergoing strong selection, it is harder to locate the favored allele in that region.

There are many directions to improve the analyses presented here. In particular, we plan to focus our attention on other organisms with more complex life cycles, experiments with variable population size and longer sampling-time-spans. As evolve and resequencing experiments continue to grow, deeper insights into adaptation will go hand in hand with improved computational analysis.

### Software and Data Availability

The source code and running scripts for Clear are publicly available at https://github.com/airanmehr/clear.

*D. melanogaster* data originally published [26, 46]. The dataset of the *D. melanogaster* study, until generation 37, is obtained from Dryad digital repository (http://datadryad.org)under accession DOI: <u>10.5061/dryad.60k68</u>. Generation 59 of the *D. melanogaster* study is accessed from European Sequence Read Archive (http://www.ebi.ac.uk/ena/) under the project accession number: PRJEB6340. The dataset containing experimental evolution of Yeast populations [14] is downloaded from http://wfitch.bio.uci.edu/~tdlong/PapersRawData/BurkeYeast.gz (last accessed 01/24/2017). UCSC browser tracks for *D. melanogaster* and Yeast data analysis are found in Suppl. Data 1 and 2, respectively.

## Acknowledgments

AI, AA, and VB were supported by grants from the NIH (1R01GM114362) and NSF (DBI-1458557 and IIS-1318386). CS is supported by the European Research Council grant ArchAdapt.

## 5 S1 Text Choosing Window Size

In genome-wide scans for detecting selection, we apply the Clear statistic on sliding windows of length *L*bp. The single locus statistic values within the window are averaged to get the composite statistic. While the statistic is robust to variation in window-size, choosing a very large window where LD has decayed will weaken the composite signal, and choosing a small window will decrease the power of composite likelihoods. Here, we use a systematic calculation to choose *L* as the distance where the LD between the favored mutation and a site *L*/2bp away remains strong.

Consider a segregating site *l* bp away from the favored allele in a selective sweep. Let *ρ*_τ_ be the LD between the favored allele and the site, τ generations after the onset of selection. Then, we have (see Eqs. 30-31 in [58]):

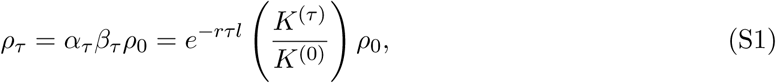

where *K*^(τ)^ = 2*ν* _τ_ (1 – *ν*_τ_) is the heterozygosity at the selected site, *r* is the recombination rate (crossovers/bp/gen). The ‘decay factor’, *α* _τ_ = *e*^−*r*τ*l*^, and ‘growth factor’, *β*_τ_, are due to recombination and selection, respectively. Under regular parameter settings, linkage to the favored allele is expected to increase after onset of selection and then decreases due to crossover events (See Fig S13-A). While *ρ*_0_ is unknown in pool-seq E&R experiments, we compute the value of *l* so that

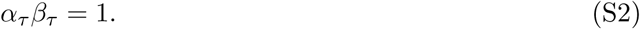

In E&R scenarios, we let τ be the time of the last sampling. For given *s*, we aim to compute the smallest window size *L* over all possible starting frequencies. Specifically,

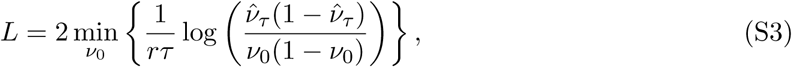

where the term *ν̂* _τ_ depends on initial frequency *ν*_0_ and selection strength *s* (Eq. 9).

We used *D. melanogaster* dataset parameters, *N* = 250, *r* = 2 × 10^−8^ and τ = 59 to compute the optimal window size for different values of *N s,* ranging from weak selection to strong selection: *N s* ∈ {20,100,200, 500}, or *s* ∈ {0.08, 0.4,0.8,2}. We set *L* = 30Kbp (See Fig S13-B) to provide good resolution for detecting weak selection.

**Fig S1:**
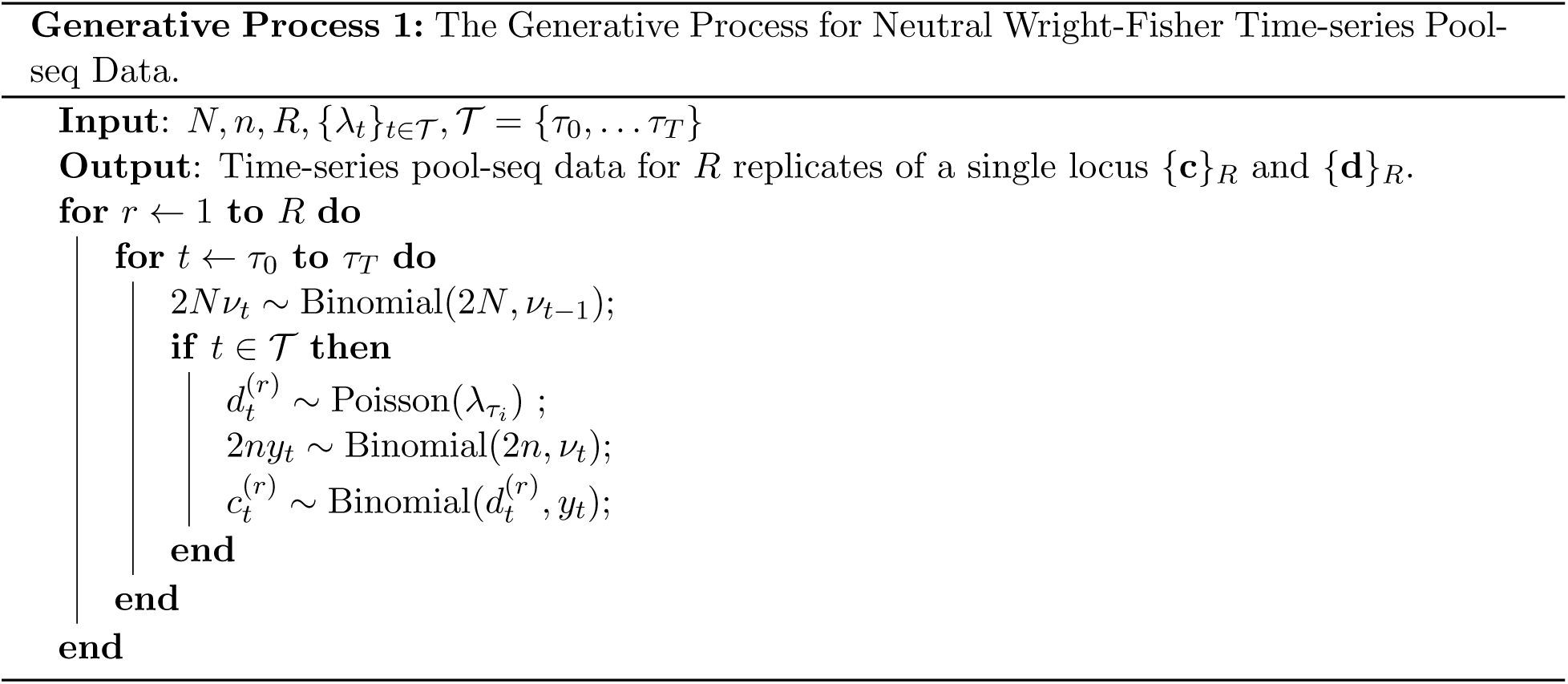
The Generative Process for Neutral Wright-Fisher Time-series Pool-seq Data.

**Fig S2:**
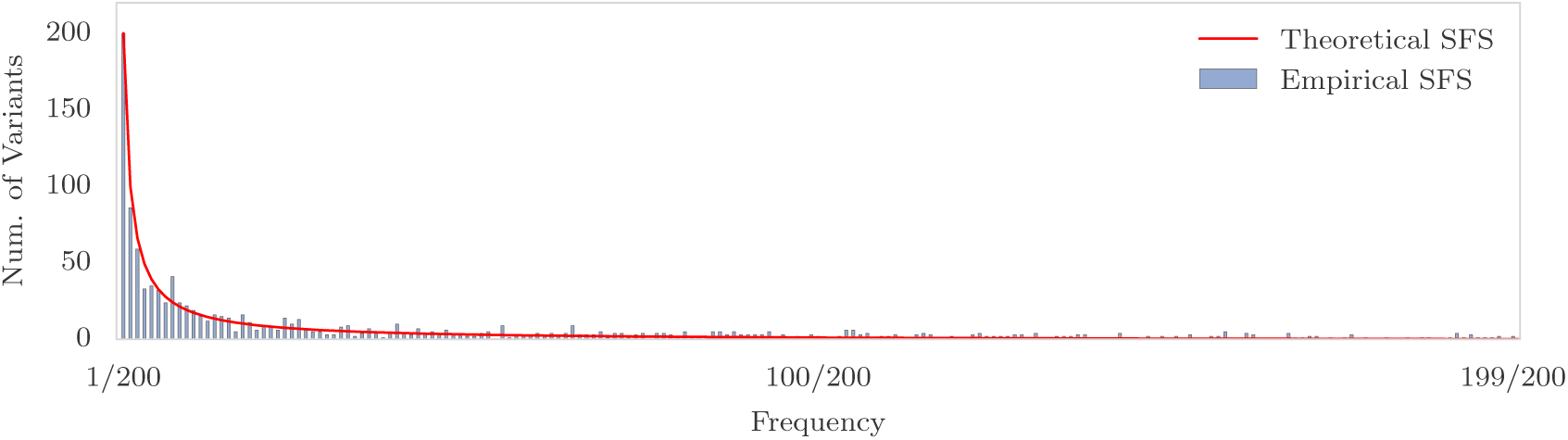
Site Frequency Spectrum. Theoretical and Empirical SFS in a 50Kbp region for a neutral population of 200 individuals when *N_e_* = 10^6^ and *μ* = 10^−9^. The *x*-axis corresponds to site frequency, and the *y*-axis to the number of variants with a specific frequency. In a neural population, majority of the variations stand in low frequency.

**Fig S3:**
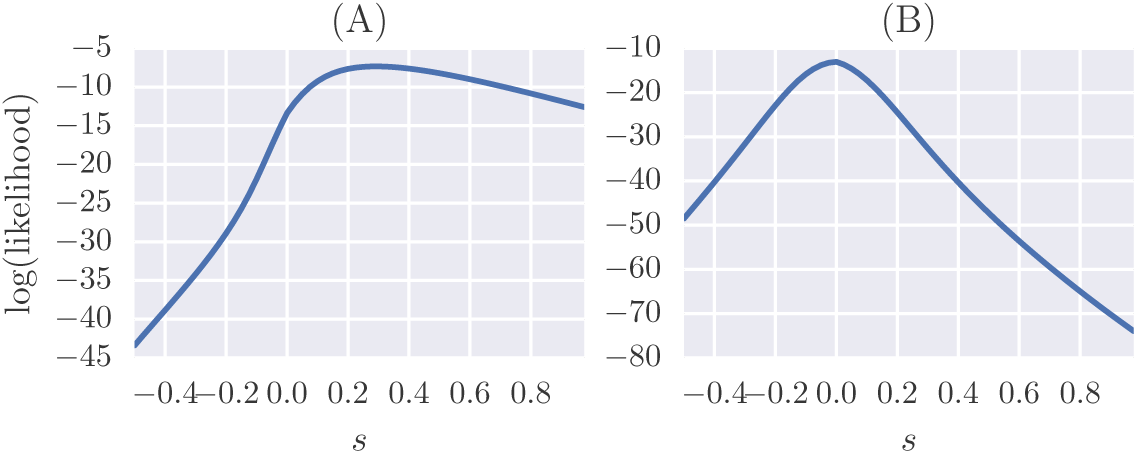
Likelihoods of the parameter *s*. Likelihood of the parameter *s* in *D. melanogaster* data for a variant with *ŝ* = 0.2 (A) and *ŝ* = 0 (B).

**Fig S4:**
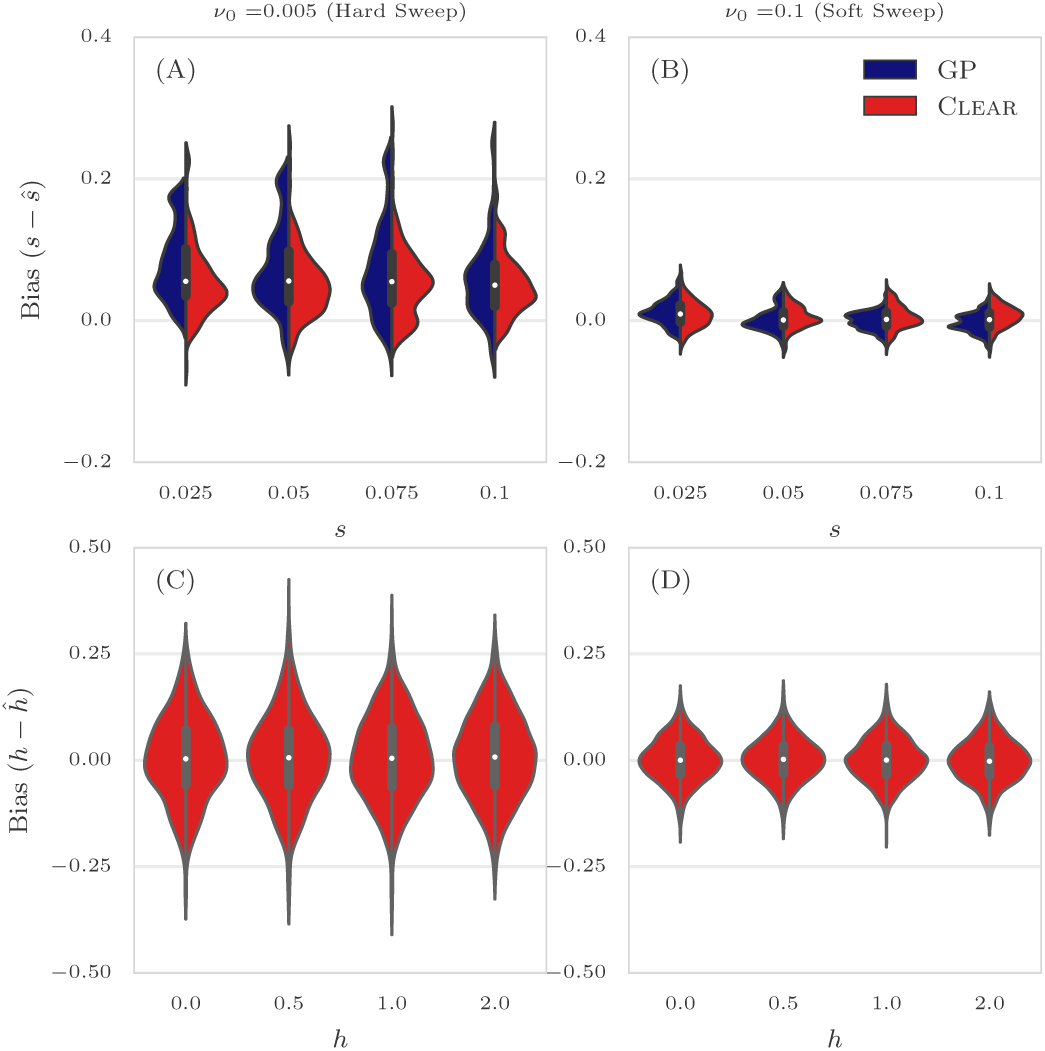
Distribution of bias for 30 × coverage. The distribution of bias (*s* – *ŝ*) in estimating selection coefficient over 1000 simulations using Gaussian Process (GP) and Clear (*H*) is shown for a range of choices for the selection coefficient s and starting carrier frequency when coverage *λ* = 30 (Panels A,B). GP and Clear have similar variance in estimates of s for soft sweep, while Clear provides lower variance in hard sweep. Also see Table S2. Panels C,D show the variance in the estimation of *h*.

**Fig S5:**
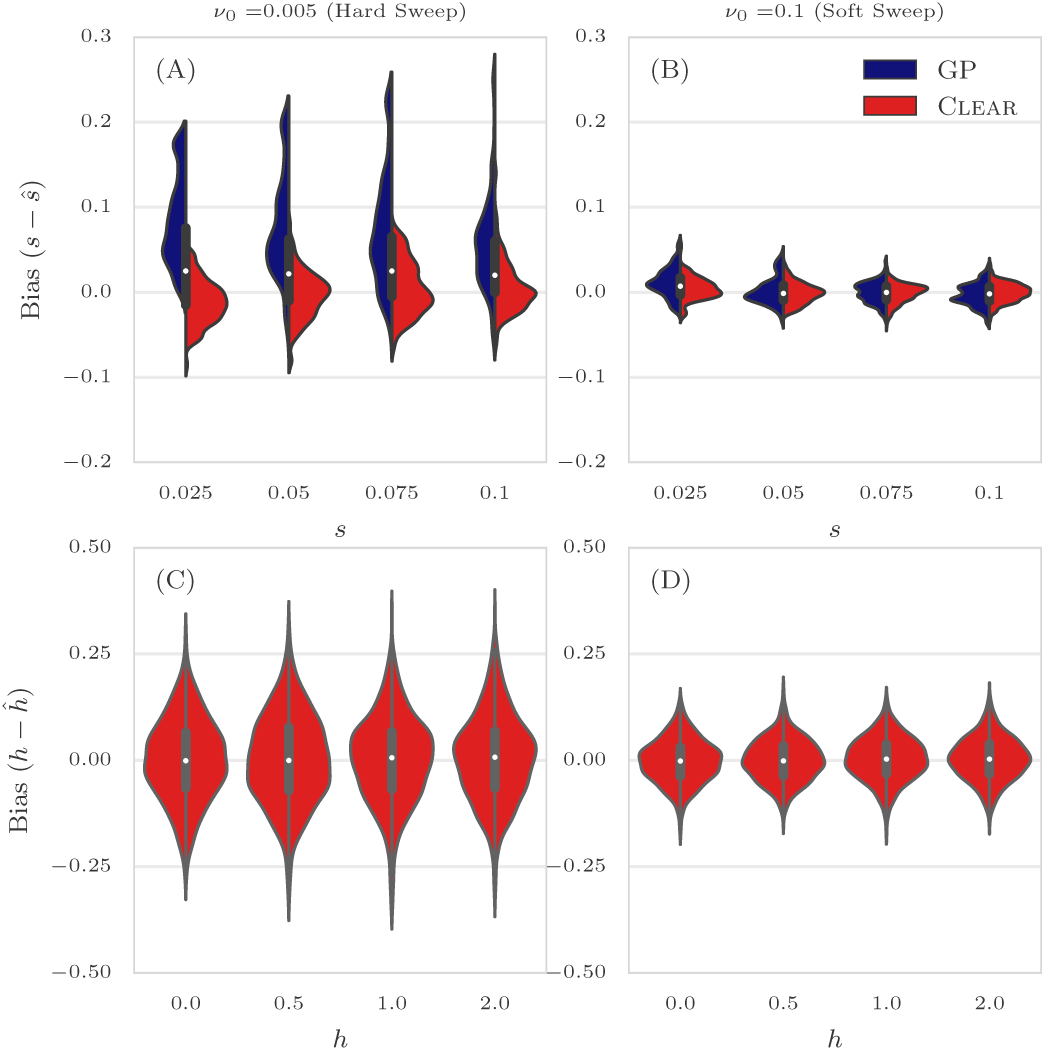
Distribution of bias for 300 × coverage. The distribution of bias (*s* – *ŝ*) in estimating selection coefficient over 1000 simulations using Gaussian Process (GP) and Clear (*H*) is shown for a range of choices for the selection coefficient s and starting carrier frequency *ν*_0_ when coverage *λ* = ∞ (Panels A,B). GP and rm better than others have similar variance in estimates of *s* for soft sweep, while Clear provides lower variance in hard sweep. Also see Table S2. Panels C,D show the variance in the estimation of *h*.

**Fig S6:**
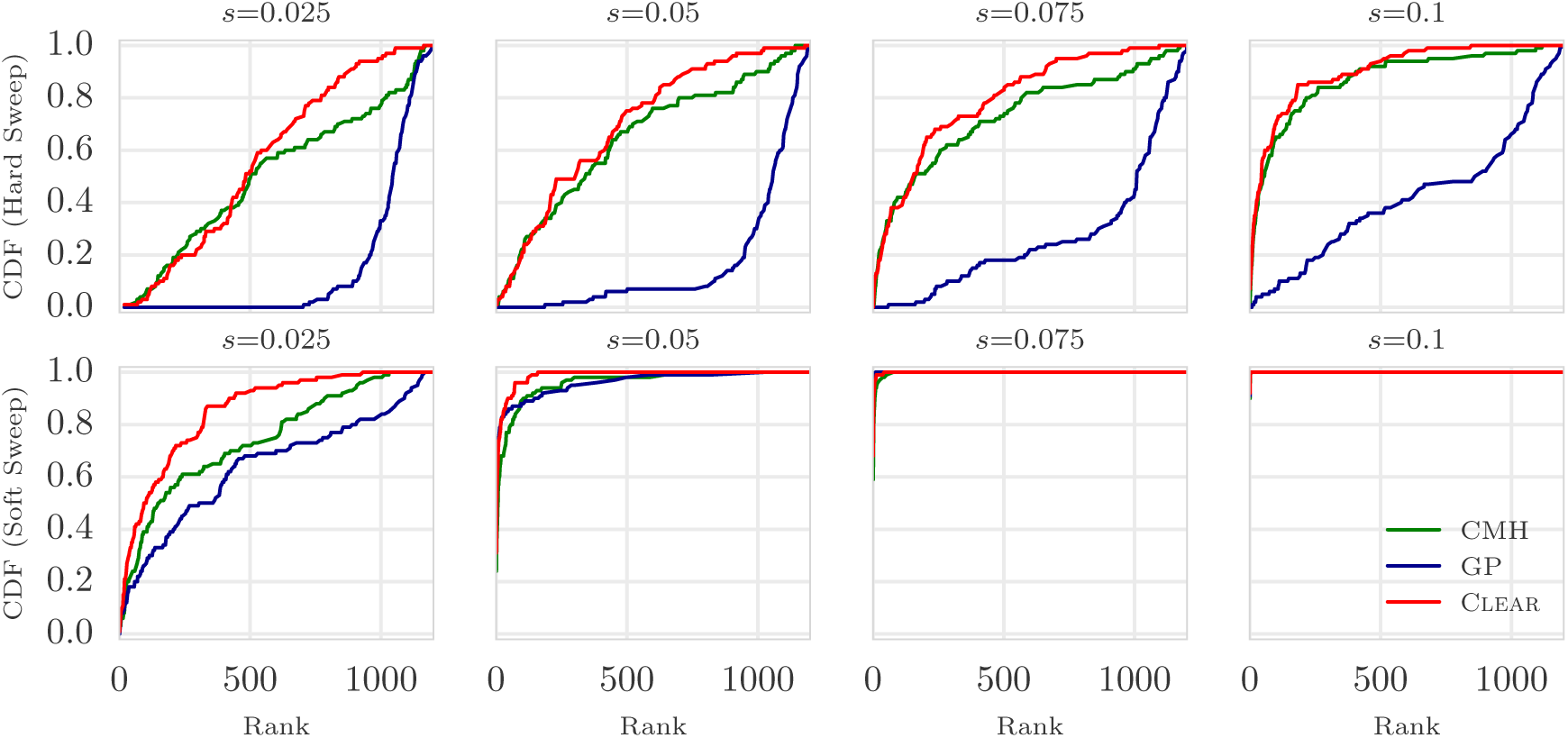
Ranking performance for 30 × coverage. Cumulative Distribution Function (CDF) of the distribution of the rank of the favored allele in 1000 simulations for Clear (*H* score), Gaussian Process (GP), and Cochran Mantel Haenszel (CMH), for different values of selection coefficient s and initial carrier frequency.

**Fig S7:**
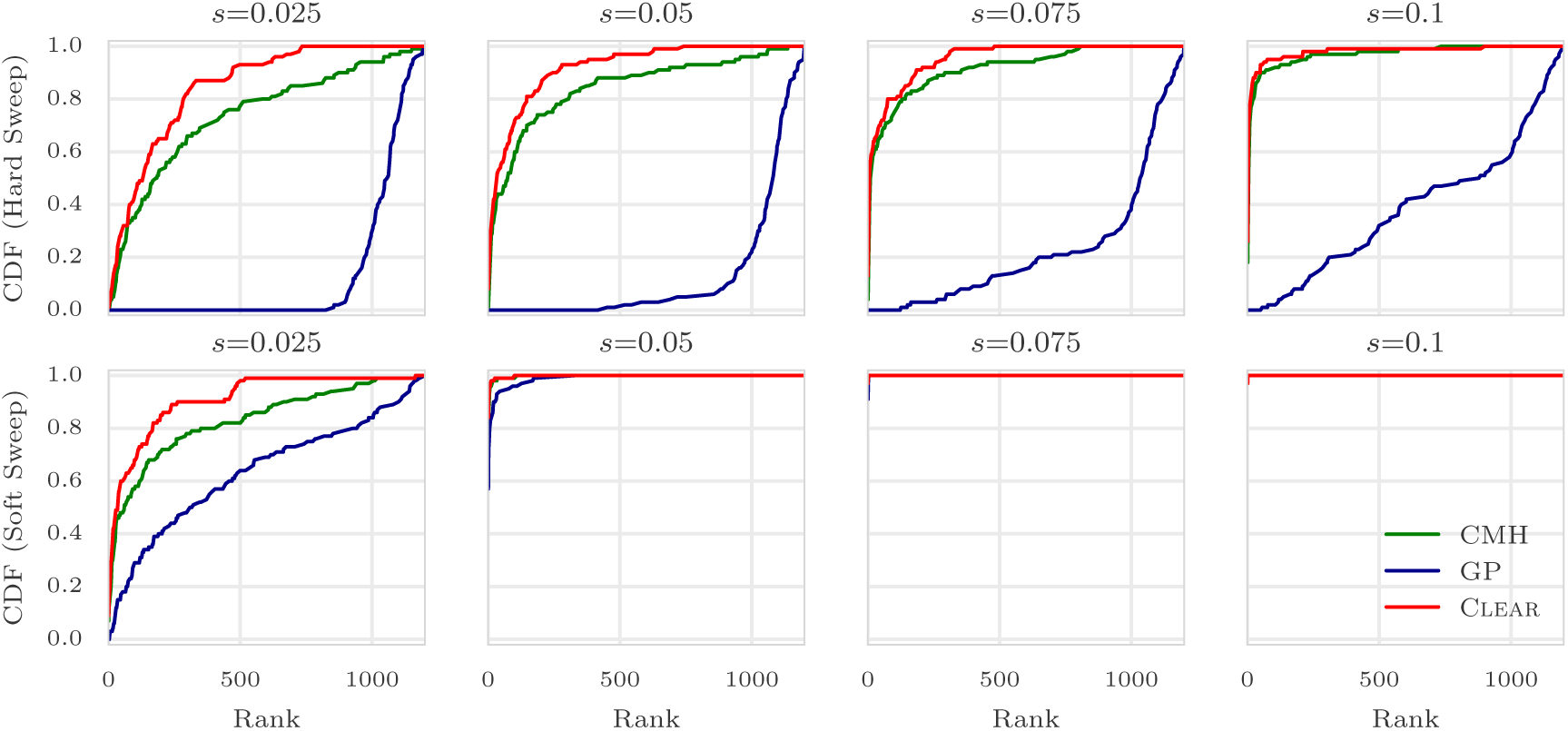
Ranking performance for 300 × coverage. Cumulative Distribution Function (CDF) of the distribution of the rank of the favored allele in 1000 simulations for Clear (*H* score), Gaussian Process (GP), and Cochran Mantel Haenszel (CMH), for different values of selection coefficient s and initial carrier frequency.

**Fig S8:**
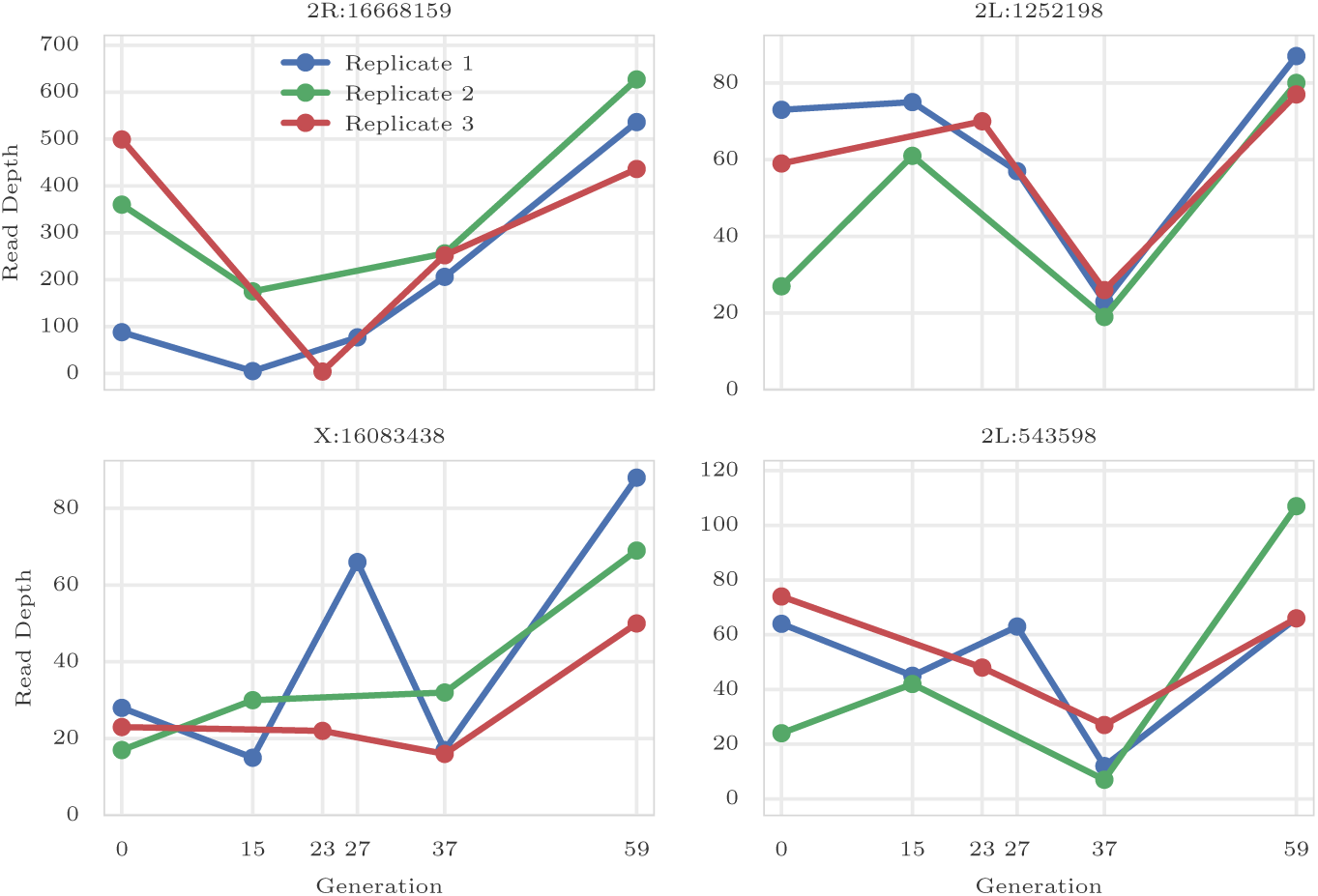
Coverage heterogeneity in time series data. Each panel shows the read depth for 3 replicates of the data from a study of *D. melanogaster* adaptation to alternating temperatures data (see section 3.1). Heterogeneity in depth of coverage is seen between replicates, and also at different time points, in all 4 variants. None of these sites pass the the hard filtering with minimum depth of 30.

**Fig S9:**
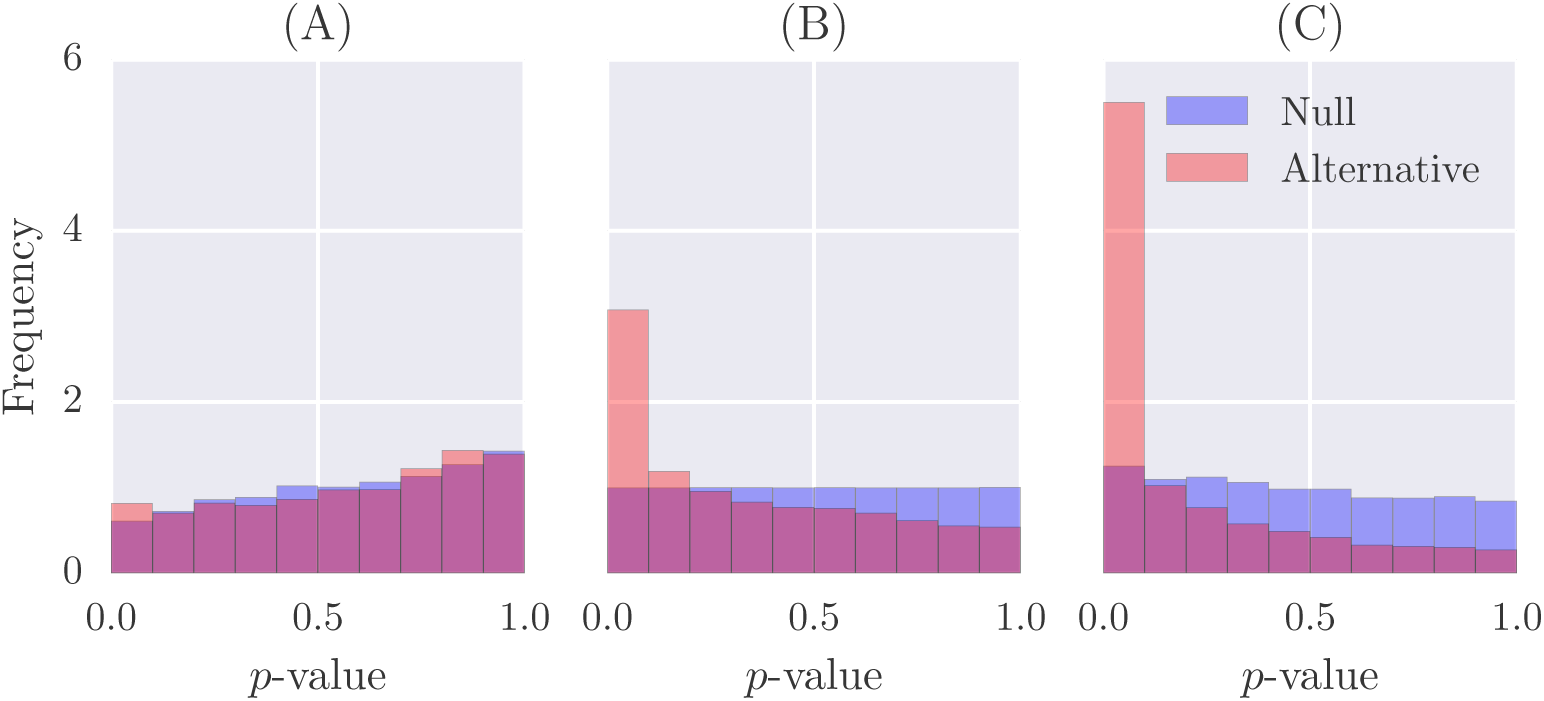
Distribution of *p*-values. Distribution of *p*-values of Clear in null simulations and experimental data when *N* = 250. Panel (A),(C) shows the effect of under estimations (*N̂* = 100) and over-estimation (*N̂* = 500) of population size in computing p-values, and panel (B) shows the distribution of *p*-values when unbiased estimate is used to create simulations.

**Fig S10:**
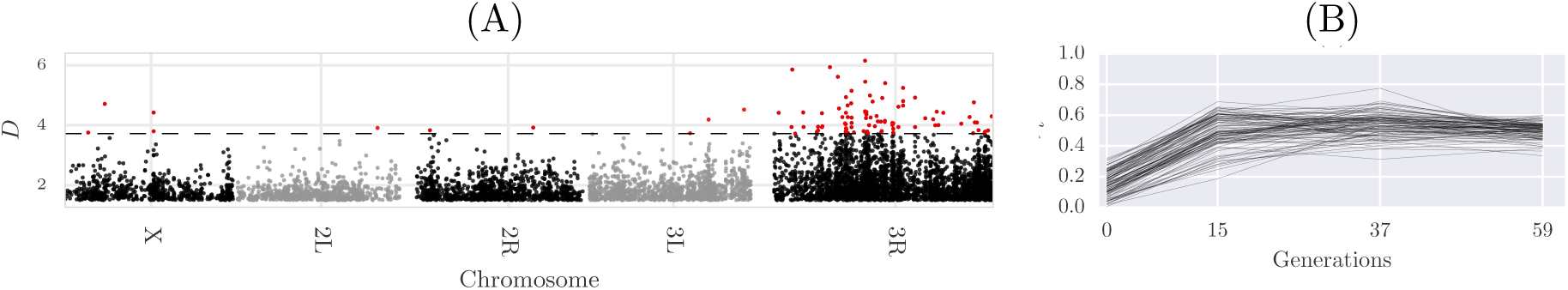
Single locus analysis of the data from a study of *D. melanogaster* adaptation to alternating temperatures. Manhattan plot of scan for testing dominant selection (A). Significant variants with FDR ≤ 0.01 are denoted in red, and their trajectories are depicted in panel (B).

**Fig S11:**
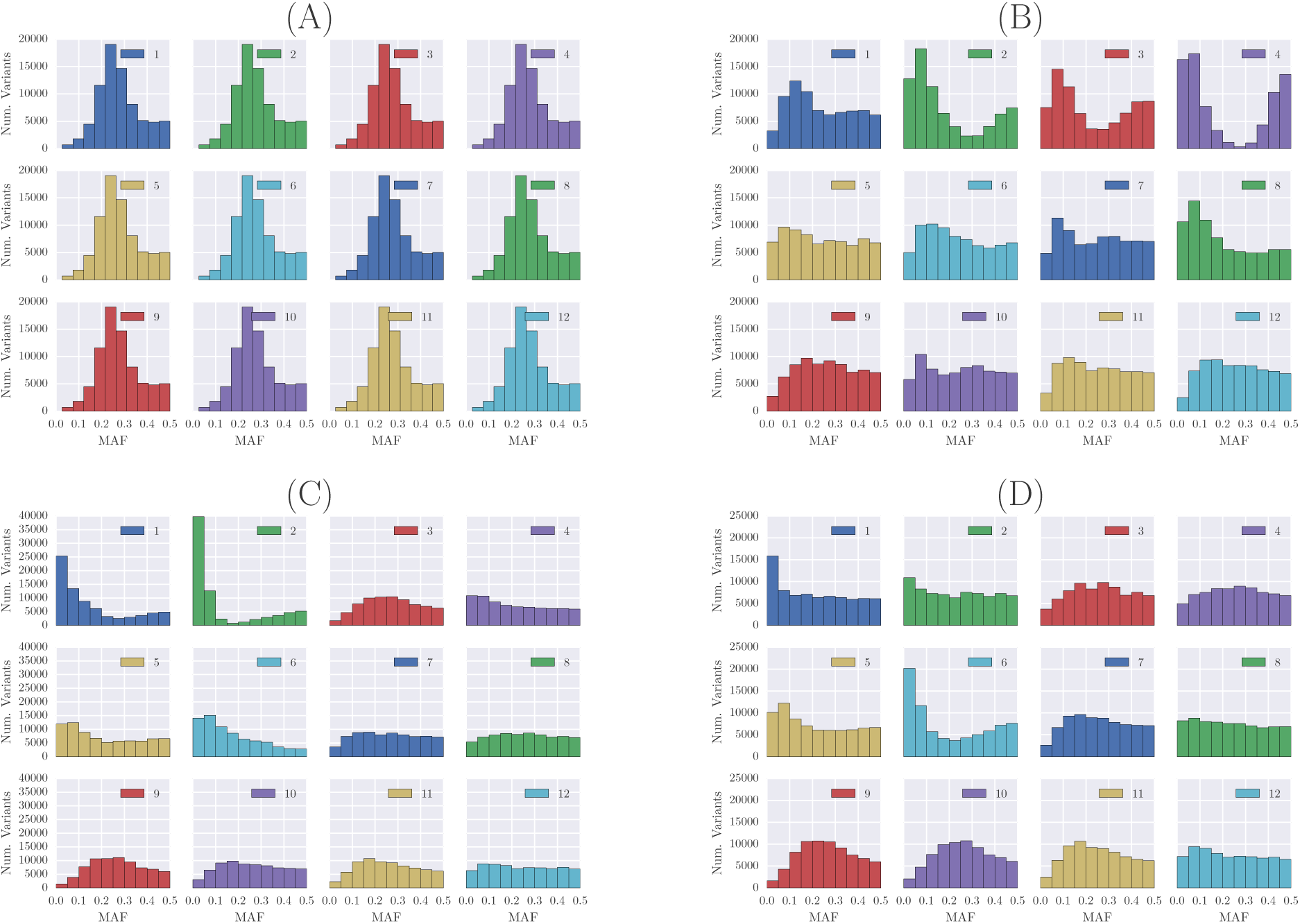
Site frequency spectrum of the Yeast dataset. Whole-genome site frequency spectrum of the Yeast dataset at generations 0 (A), 180 (B), 360 (C) and 540 (D). Some replicates, e.g. replicate 2, undergoing severe demographic events.

**Fig S12:**
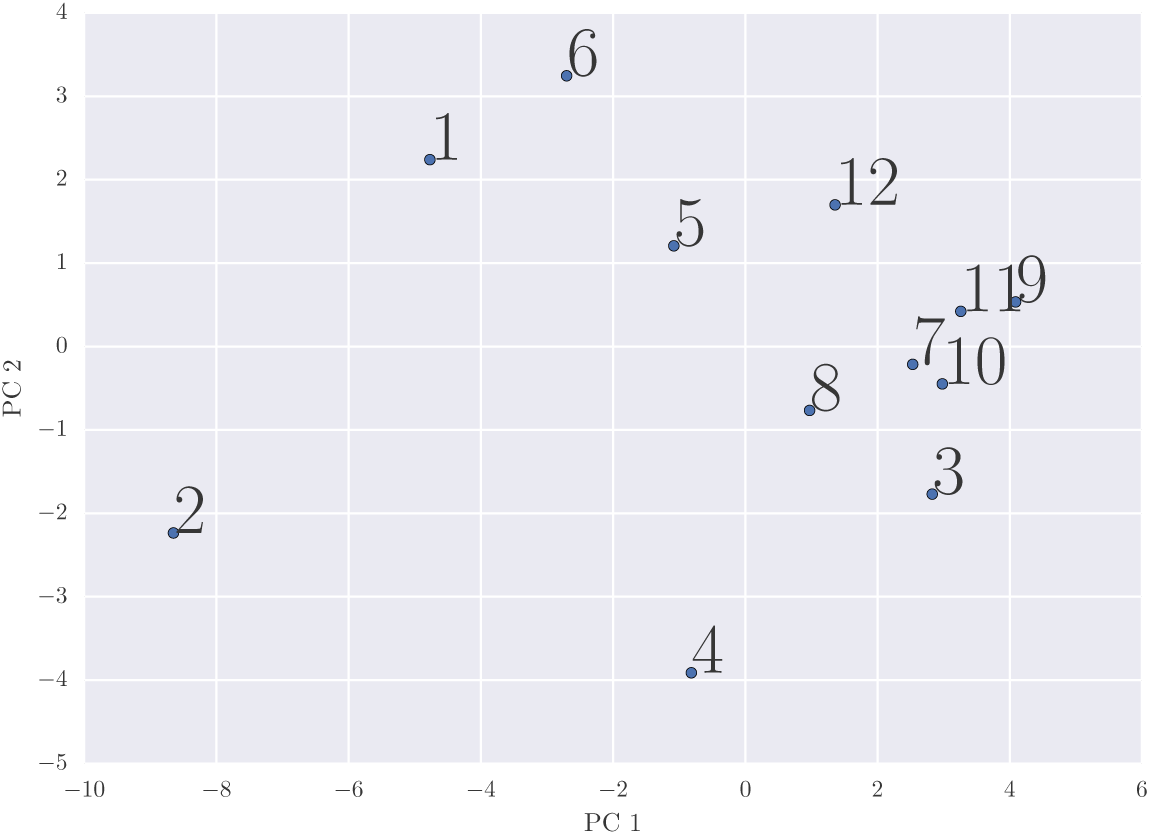
Population similarity. Principle component analysis of the 12 replicates throughout the experiment, showing that some populations exhibiting distinct frequency spectra.

**Fig S13:**
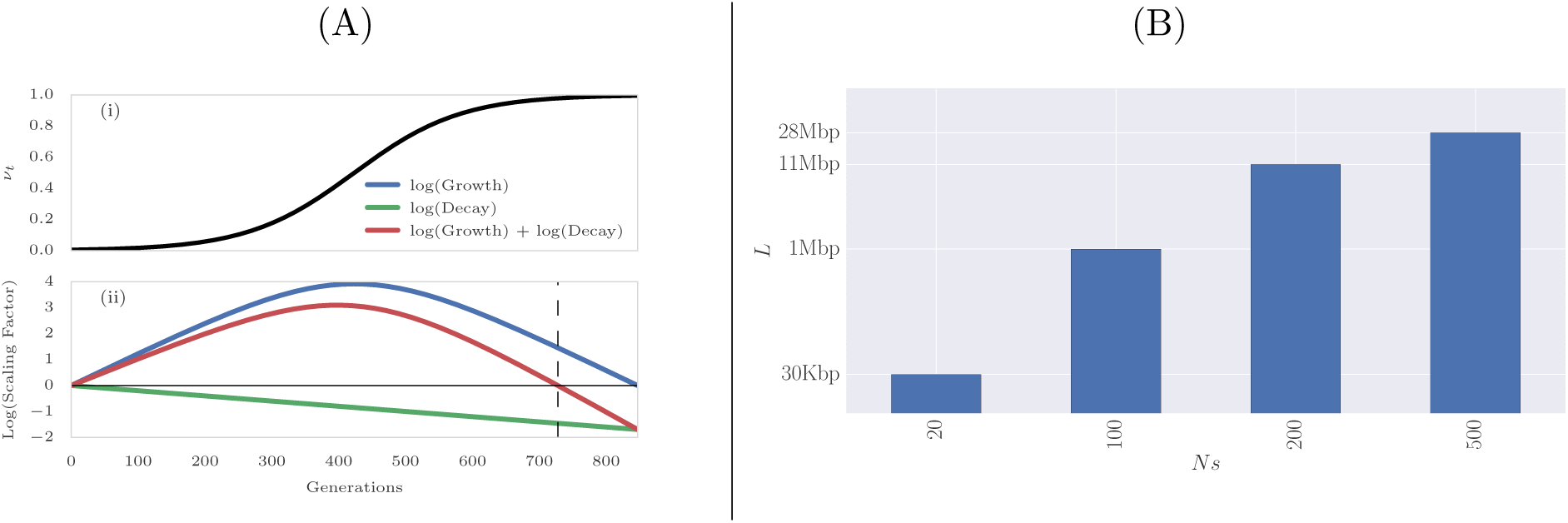
Choosing window size for Clear statistic. (A) Expected dynamics of LD between favored allele (*s* = 0.025) and a variant 50Kbp away, with initial frequency *ν*_*0*_ = 0.01. (A-i) depicts the dynamic of the favored allele during the selective sweep. (A-ii) illustrates interaction of the growth and decay factors introduced in Eq. S1, with the red line describing overall effect of selection and recombination on LD. The vertical dashed line points to the time when the LD value starts to decrease below original LD. (B) Alternatively, we can fix time, and find the window-size at which LD decays below the original LD (Eq. S3). The plot shows the window size as a function of *N s* (20,100,200,500), after fixing other model parameters to match *D. melanogaster* E&R experiments (*N* = 250, *r* = 2 × 10^8^, τ= 59).

**Table S1:** Average of power for detecting selection. Average power is computed for 8000 simulations with *s* ∈ {0.025,0.05,0.075,0.1}. Frequency Increment Test (FIT), Gaussian Process (GP), Clear (𝓗 statistic) and Cochran Mantel Haenszel (CMH) are compared for different initial carrier frequency *ν*_0_. For all sequencing coverages, Clear outperform other methods. When coverage is not high (*λ* ∈ {30,100}) and initial frequency is low (hard sweep), Clear significantly perform better than others.

| Hard Sweep |  |  | Soft Sweep |  |  |
| --- | --- | --- | --- | --- | --- |
| $\lambda$ | Method | Avg Power | $\lambda$ | Method | Avg Power |
| 300 | CLEAR | 34 | 300 | CLEAR | 69 |
| 300 | CMH | 12 | 300 | CMH | 69 |
| 300 | FIT | 2 | 300 | GP | 61 |
| 300 | GP | 0 | 300 | FIT | 8 |
| 100 | CLEAR | 31 | 100 | CLEAR | 67 |
| 100 | CMH | 4 | 100 | CMH | 60 |
| 100 | FIT | 2 | 100 | GP | 59 |
| 100 | GP | 0 | 100 | FIT | 1 |
| 30 | CLEAR | 20 | 30 | CLEAR | 57 |
| 30 | FIT | 2 | 30 | GP | 53 |
| 30 | CMH | 0 | 30 | CMH | 39 |
| 30 | GP | 0 | 30 | FIT | 3 |

**Table S2:**
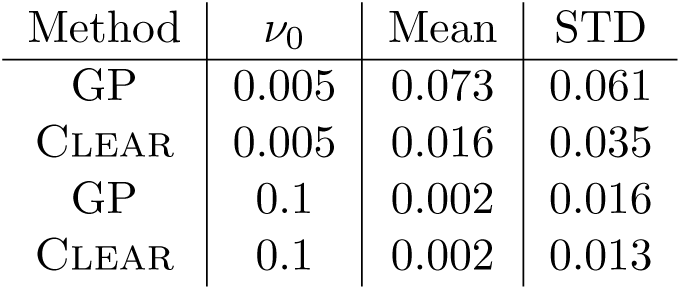
Mean and standard deviation of the distribution of bias (*s* – *ŝ*) of 8000 simulations with coverage *λ* = 100× **and** *s* ∈ {0.025, 0.05, 0.075, 0.1}.

| Method | $\nu_0$ | Mean | STD |
| --- | --- | --- | --- |
| GP | 0.005 | 0.073 | 0.061 |
| CLEAR | 0.005 | 0.016 | 0.035 |
| GP | 0.1 | 0.002 | 0.016 |
| CLEAR | 0.1 | 0.002 | 0.013 |

**Table S3:**
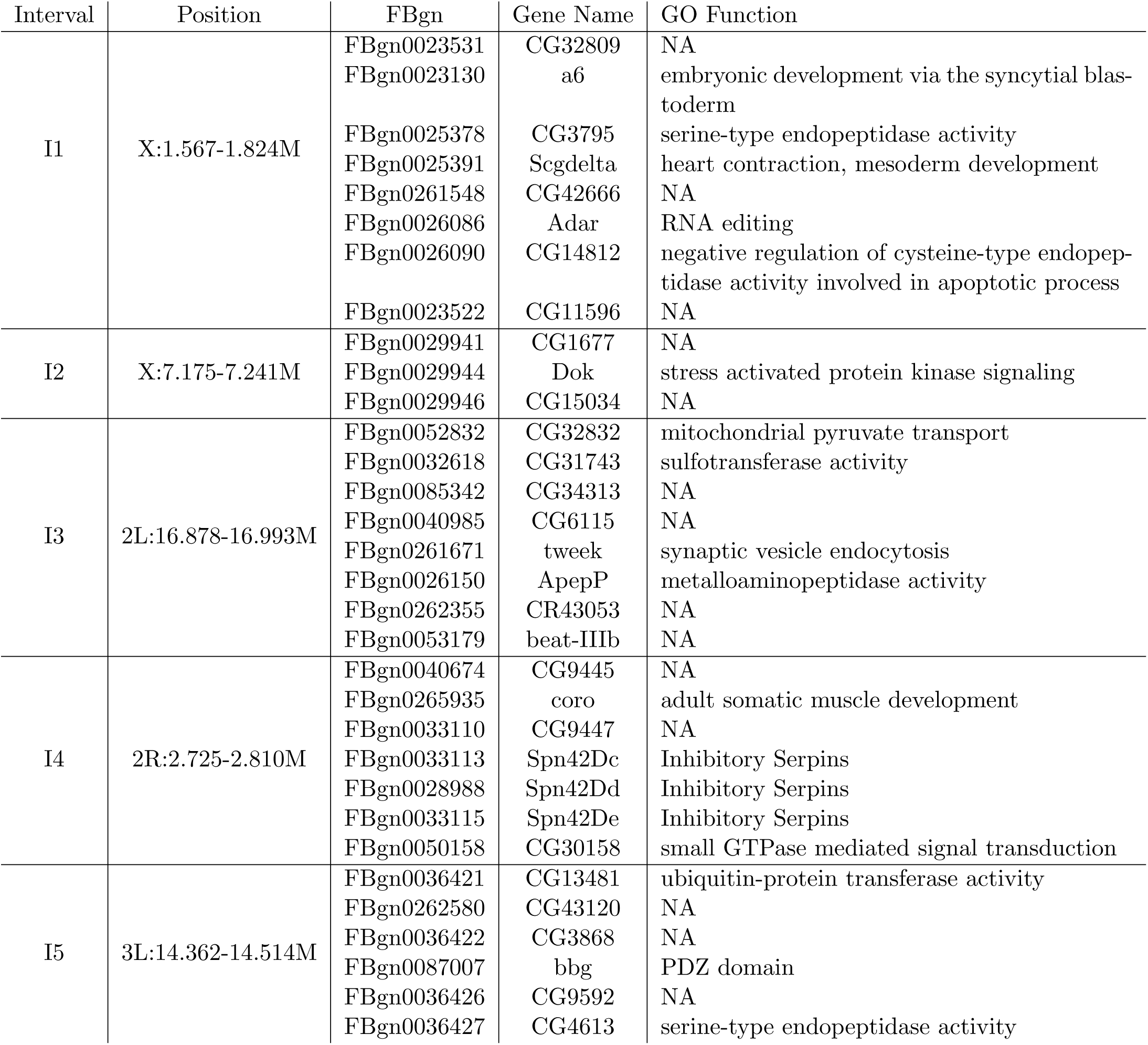
Overlapping genes with the 174 candidate variants.

